# Novel patterns of complex structural variation revealed across thousands of cancer genome graphs

**DOI:** 10.1101/836296

**Authors:** Kevin Hadi, Xiaotong Yao, Julie M. Behr, Aditya Deshpande, Charalampos Xanthopoulakis, Joel Rosiene, Madison Darmofal, Huasong Tian, Joseph DeRose, Rick Mortensen, Emily M. Adney, Zoran Gajic, Kenneth Eng, Jeremiah A. Wala, Kazimierz O. Wrzeszczyński, Kanika Arora, Minita Shah, Anne-Katrin Emde, Vanessa Felice, Mayu O. Frank, Robert B. Darnell, Mahmoud Ghandi, Franklin Huang, John Maciejowski, Titia De Lange, Jeremy Setton, Nadeem Riaz, Jorge S. Reis-Filho, Simon Powell, David Knowles, Ed Reznik, Bud Mishra, Rameen Beroukhim, Michael C. Zody, Nicolas Robine, Kenji M. Oman, Carissa A. Sanchez, Mary K. Kuhner, Lucian P. Smith, Patricia C. Galipeau, Thomas G. Paulson, Brian J. Reid, Xiaohong Li, David Wilkes, Andrea Sboner, Juan Miguel Mosquera, Olivier Elemento, Marcin Imielinski

**Author notes:** These authors contributed equally (Kevin Hadi, Xiaotong Yao, Julie M. Behr).

## Abstract

Cancer genomes often harbor hundreds of somatic DNA rearrangement junctions, many of which cannot be easily classified into simple (e.g. deletion, translocation) or complex (e.g. chromothripsis, chromoplexy) structural variant classes. Applying a novel genome graph computational paradigm to analyze the topology of junction copy number (JCN) across 2,833 tumor whole genome sequences (WGS), we introduce three complex rearrangement phenomena: *pyrgo, rigma*, and *tyfonas*. Pyrgo are “towers” of low-JCN duplications associated with early replicating regions and superenhancers, and are enriched in breast and ovarian cancers. Rigma comprise “chasms” of low-JCN deletions at late-replicating fragile sites in esophageal and other gastrointestinal (GI) adenocarcinomas. Tyfonas are “typhoons” of high-JCN junctions and fold back inversions that are enriched in acral but not cutaneous melanoma and associated with a previously uncharacterized mutational process of non-APOBEC kataegis. Clustering of tumors according to genome graph-derived features identifies subgroups associated with DNA repair defects and poor prognosis.

## Introduction

Cancer genomes are shaped by both simple and complex structural DNA variants (SV). While simple structural variants (e.g. deletions, duplications, translocations, inversions) arise through the breakage and fusion of a few (1-2) genomic locations, complex SVs can cause multiple (≥ 2) DNA junctions harboring distinct reference genome topologies and yield one or more copies of complex rearranged alleles (Maciejowski and Imielinski, 2017). Though many mechanisms have been postulated to explain complex SV patterns (chromothripsis (Stephens et al., 2011), chromoplexy (Baca et al., 2013), templated insertion chains (TIC), (Li et al., 2017; Spies et al., 2017), breakage fusion bridge cycles (Zakov et al., 2013)), the field has not yet converged to a single unifying framework to identify these patterns in a tumor whole genome sequence. In addition, it is unclear whether some of the clustered rearrangement patterns commonly observed in cancer represent as yet uncharacterized event classes. As a result, the mutational processes driving complex SV evolution are still obscure.

Though the detection of variant DNA junctions (two reference genome locations with orientations joined in one of four basic orientations: deletion-like (DEL-like), tandem duplication-like (DUP-like), inversion-like (INV-like), translocation-like (TRA-like, **Fig. S1B**) in WGS is routine, the classification of junctions into events (e.g. deletion, duplication, inversion, translocation) becomes difficult when two or more junctions are near each other. Furthermore, rearrangements and copy number alterations (CNA) are usually analyzed and interpreted separately, despite being two facets of a single genome structure.

In WGS, CNAs are detected as change-points (Chiang et al., 2009) in sequencing read depth along the genome (e.g. BIC-seq (Xi et al., 2011)) while rearrangement junctions are nominated through the analysis of junction-spanning read pairs (e.g. SvABA (Wala et al., 2018), GRIDSS (Cameron et al., 2017)). CNAs and junctions are, however, intrinsically coupled, since every copy of every non-telomeric segment must have a left (towards smaller coordinates) and a right (towards larger coordinates) neighbor, whether that neighbor is already adjacent on the reference (via reference or REF junction) or introduced through rearrangement (via a variant or ALT junction) (Medvedev et al., 2010; Greenman et al., 2012). Furthermore, though copy number (CN) is a concept primarily applied to describe the dosage of genomic *intervals*, a DNA junction may also be present in one or more copies per cell, and thus be assigned a junction copy number (JCN), i.e. the number of alleles harboring the rearrangement at a given locus. We hypothesized that the topology and dosage of both intervals and junctions on genome graphs can provide a source of important features to classify complex SVs and define novel mutational processes. To address this hypothesis, we assembled a dataset of nearly 3,000 WGS cases spanning 31 tumor types.

## Results

### JaBbA accurately infers junction-balanced genome graphs

We developed an algorithm (Junction Balance Analysis, JaBbA) to investigate the topology of junction copy number in cancer genomes. JaBbA infers *junction-balanced genome graphs* (see Methods for detailed formulation). We define a genome graph as a directed graph whose vertices are strands of genomic segments and whose directed edges each represent a pair of 3’ and 5’ DNA ends that are adjacent in the reference (REF edge) or connected through rearrangement (ALT edge). A *junction balanced genome-graph* assigns every graph node and edge an integer copy number, while enforcing the constraint that every copy of every interval must have a left and a right neighbor (i.e. in the reference genome, or introduced through rearrangement).

JaBbA takes normalized read depth (across 200 bp bins) and junctions (e.g. nominated by SvABA) as input, and applies a probabilistic model to minimize the residual between observed read depth and inferred interval dosage through joint assignment of copy number to intervals and junctions (**Fig. 1A**). The resulting junction-balanced genome graph obeys the network constraints that the dosage of every interval (vertex) is equal to the sum of the copy numbers of incoming (similarly, outgoing) junctions (**Fig. S1A**). Since short-read WGS may fail to detect certain junctions (e.g. those connecting one or two low mappability breakpoints), we allow the model to harbor occasional *loose ends*, which represent CN change-points that cannot be associated with a nearby junction. By penalizing loose ends within the objective function of the model, JaBbA achieves an optimal fit of integer copy numbers genome wide to both segments and junctions, while accounting for incomplete input data.

**Fig. 1.**
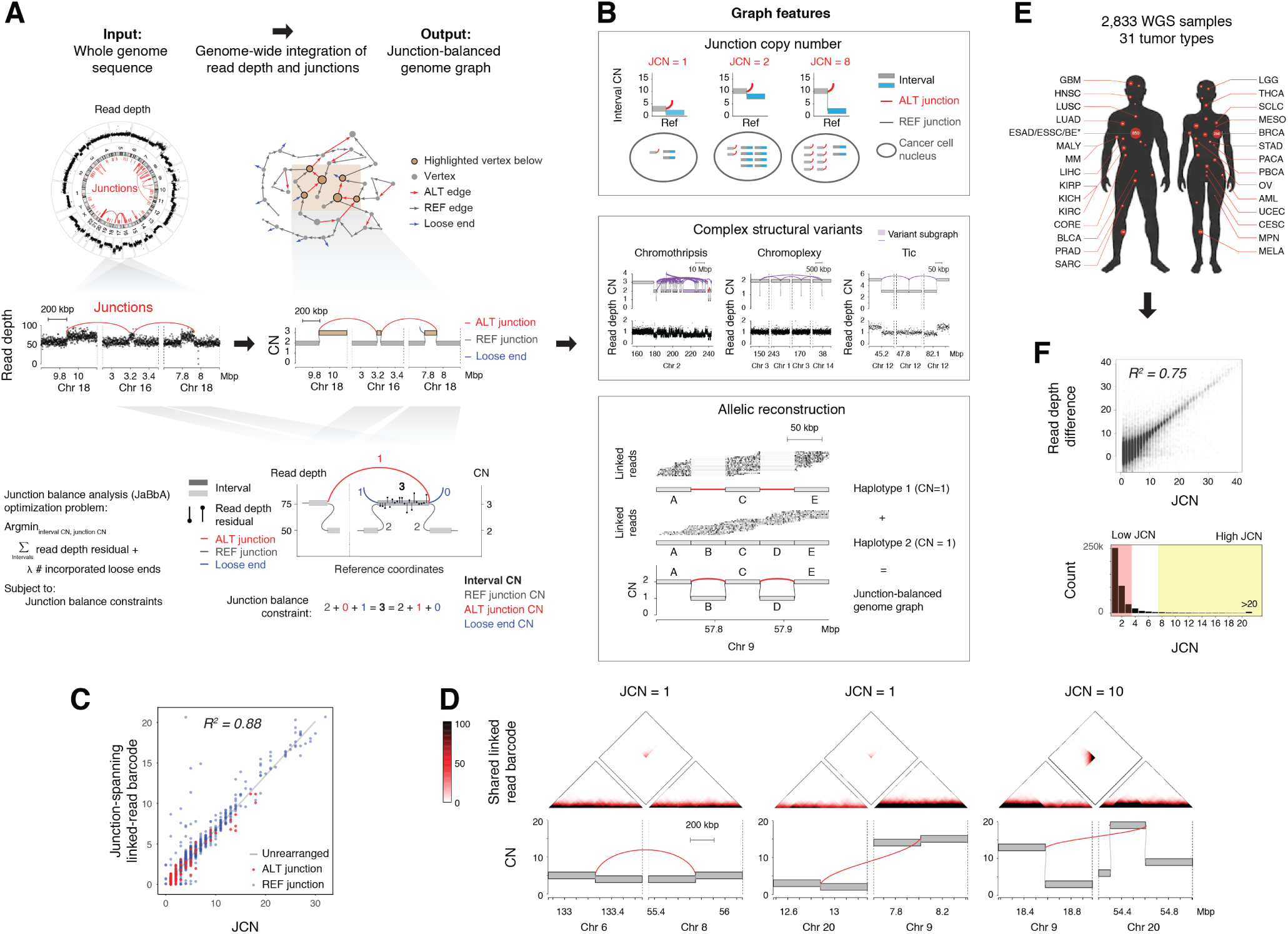
Junction-balanced graphs as a novel paradigm for the integrative analysis of rearranged genomes. (A) Junction balance analysis (JaBbA) integrates high density WGS read depth data and rearrangement junctions to estimate junction copy number (JCN) and generates coherent models of genome structure. Bottom, a schematic of the mixed integer quadratic program optimization problem which JaBbA solves. (B) Selected applications of JaBbA graphs. Top, JCN refers to the number of copies per cell of a junction. Middle, detection of known and novel complex rearrangement events as subgraphs through the analysis of graph features. Bottom panel, using segmental and junction copy number, feasible reconstructions of allelic haplotypes can be inferred through the deconvolution of JaBbA graphs, which when summed give rise to the observed JaBbA graph. 10X linked-read sequencing (top and middle tracks) can be used to constrain these reconstructions. (C) The number of junction-spanning 10X linked-read barcodes within rearranged loci as detected by JaBbA on WGS data shows high correlation to the estimated JCN. (D) Top, heatmap of the number of shared 10X linked reads between the pair of genomic locations. Bottom, JaBbA output genome graph within 20 kbp of the featured junctions. (E) Cohort of 2,833 tumor/normal pairs across 2,552 patients and 31 tumor types; abbreviations can be found in **Table 1**. ESAD and BE have patients with multiple samples. (F) Summary of fitted JCN. Top, purity and ploidy corrected read depth difference between the *cis* and *trans* side of breakpoints over the JaBbA-fitted JCN. Bottom, histogram of JCNs in the cohort. Other abbreviations: CN, copy number; Tic, templated insertion chain.

In our benchmarking experiments (see Methods), JaBbA inferred JCN with consistently higher fidelity than previously published genome graph-based methods (ReMixT (McPherson et al., 2017), Weaver (Li et al., 2016), PREGO (Oesper et al., 2012)) across a wide range of tumor purities (**Fig. S1C-D**). In addition, JaBbA consistently outperformed classic (i.e. non-graph based) CNA callers (BIC-seq (Xi et al., 2011), FACETS (Shen and Seshan, 2016), TITAN (Ha et al., 2014), FREEC (Boeva et al., 2012), CONSERTING (Chen et al., 2015)) and genome graph-based methods in estimating interval CN amplitude and CN change-point locations across a wide range of tumor purities (**Fig. S1C-F**). This also resulted in closer visual correspondence between fitted graphs and observed read depth patterns (**Fig. S1G**) and closer colocalization between rearrangement junctions and CN change-points (**Fig. S1H**). We attribute JaBbA’s superior performance to its use of a loose end prior for regularization as well as a global optimization (mixed integer program) rather than a local optimization (expectation maximization for ReMixT, loopy belief propagation for Weaver) algorithm for model fitting.

**Table 1.** Tumor types collected and analyzed in this study.

| Abbr. | Tumor type | N |
| --- | --- | --- |
| AML | acute myeloid leukemia | 56 |
| BE | Barrett's esophagus | 347 |
| BLCA | bladder urothelial carcinoma | 38 |
| BRCA | breast invasive carcinoma | 262 |
| CESC | cervical squamous cell carcinoma and endocervical adenocarcinoma | 20 |
| CORE | colon or rectum adenocarcinoma | 106 |
| ESAD | esophageal adenocarcinoma | 432 |
| ESSC | esophageal squamous cell carcinoma | 20 |
| GBM | glioblastoma multiforme | 83 |
| HNSC | head-and-neck squamous cell carcinoma | 61 |
| KICH | chromophobe renal cell carcinoma | 51 |
| KIRC | kidney renal clear cell carcinoma | 62 |
| KIRP | kidney renal papillary cell carcinoma | 41 |
| LGG | lower grade glioma | 57 |
| LIHC | liver hepatocellular carcinoma | 66 |
| LUAD | lung adenocarcinoma | 149 |
| LUSC | lung squamous cell carcinoma | 68 |
| MALY | malignant lymphoma | 111 |
| MELA | melanoma | 256 |
| MESO | mesothelioma | 4 |
| MM | multiple myeloma | 4 |
| MPN | myeloproliferative neoplasms | 1 |
| OV | ovarian serous cystadenocarcinoma | 72 |
| PACA | pancreatic carcinoma | 27 |
| PBCA | pediatric brain cancer | 4 |
| PRAD | prostate adenocarcinoma | 115 |
| SARC | sarcoma | 57 |
| SCLC | small cell lung cancer | 46 |
| STAD | stomach adenocarcinoma | 64 |
| THCA | thyroid carcinoma | 62 |
| UCEC | uterine corpus endometrial carcinoma | 64 |
| OTHER |  | 27 |

We orthogonally validated JaBbA’s ability to infer JCN from short-read WGS with 10X Chromium linked-read WGS (**Fig. 1C-D**). In the breast cancer cell line HCC1954, JaBbA-derived JCN estimates closely correlated with the read density of junction-spanning linked-read barcodes (*R*^2^ = 0.88) (**Fig. 1C**). This included low copy (JCN=1) junctions connecting both low copy (CN < ploidy) and high copy (CN=14) intervals, as well as high copy (CN=10) junctions (**Fig. 1D**). These results show that JCN is a property that can be robustly inferred from short read WGS and is independent from interval CN.

### Pan-cancer analysis of junction-balanced genome graphs

To investigate the topology of junction copy number across cancer, we assembled a dataset comprising 2,833 short-read WGS tumor or cell line samples spanning 31 primary tumor types (**Fig. 1E, Table 1**). Among these, we generated WGS for 546 previously unpublished WGS tumor/normal pairs, including 199 precision oncology cases from 6 New York City-based cancer centers and 347 pre-malignant Barrett’s esophagus and gastric tumors from 80 patients (Paulson et al., 2019) **Fig. 1E, Table S1** (A companion study investigating Barrett’s esophagus WGS dataset in detail is currently in preparation). In summary, our analysis included 1,668 WGS samples not currently within the Pan-Cancer Analysis of Whole Genomes (PCAWG) effort, including published studies (Nik-Zainal et al., 2016; Hayward et al., 2017; Baca et al., 2013; Lee et al., 2019; Barretina et al., 2012; Frankell et al., 2019), The Cancer Genome Atlas (TCGA, 1017 cases), International Cancer Genome Consortium (ICGC, 876 cases), or Cancer Cell Line Encyclopedia (CCLE, 326 cases) (Barretina et al., 2012; Ghandi et al., 2019). Though the majority of our analyzed samples consisted of primary tumors, 283 out of the 2,833 samples were extracted or derived from metastatic tumors.

Application of harmonized pipelines for high-density read depth calculation and junction calling (SvaBA) followed by JaBbA (**Fig. 1A-B**) to these 2,833 samples yielded 2,798 high quality genome graphs (see Methods for quality control and reasons for sample exclusion, also **Fig. S5B**). Analyzing junction-balanced genome graph topology, we identified subgraphs associated with previously identified complex rearrangement patterns such as chromothripsis, chromoplexy, and TICs (**Fig. 1B, middle**) implementing criteria described in previous publications within our framework (see Methods). Consistent with our 10X Chromium WGS benchmarks (see above) (**Fig. 1C**), we observed wide variation in inferred JCN across our datasets which correlated with observed read depth changes at junction breakpoints. While the vast majority of junctions demonstrated low-JCN (JCN < 4), we observed a long tail of high-JCN (JCN > 7) junctions (**Fig. 1F**).

### Low-JCN junctions cluster into towers and chasms

To distinguish between complex SV patterns associated with low-JCN vs. high-JCN junctions, we identified junction clusters based on their overlapping footprints on the reference and labeled each cluster as high- / low- JCN on the basis of its highest copy junction. Considering clusters harboring three or more junctions, we found that low-JCN clusters were more likely to be dominated by a single junction type (> 90% representation of that type). Specifically, we found low-JCN clusters were significantly more likely to be predominantly composed of DEL-like (*P* < 2.2 × 10^−16^, z-test, logistic regression) or DUP-like junctions (*P* < 2.2 × 10^−16^) **Fig. S2A**).

To rigorously nominate clusters of low copy DUP-like and DEL-like junctions in each tumor sample, we identified genomic genomic bins (1 Mbp width, 500 kbp stride) harboring more low-JCN of a given type (e.g. DEL-like) than expected under a gamma-Poisson background model, employing the total count of other (e.g. non-DEL like) junction classes as a covariate. We found excellent model fits for both DUP-like (**Fig. 2A**) and DEL-like (**Fig. 2B**) analyses, as shown by quantile-quantile (Q-Q) plots (genomic inflation factor, *λ*, near 1) harboring a set of significant outliers. In each analysis, non-outlier data points comprised bins harboring visually apparent simple deletions or duplications (**Fig. 2A-B**). Outlier bins in the DUP-like model corresponded to “towers” of low-JCN duplications (**Fig. 2A**; **Fig. S2C**) that we named *pyrgo* (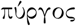, Greek meaning tower). Outlier loci in the DEL-like models comprised subgraphs of interval CN “chasms” flanked by low-JCN deletions whose interval CN often reached 0 (**Fig. 2B**). We named these patterns *rigma* (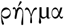, Greek meaning rift).

**Fig. 2.**
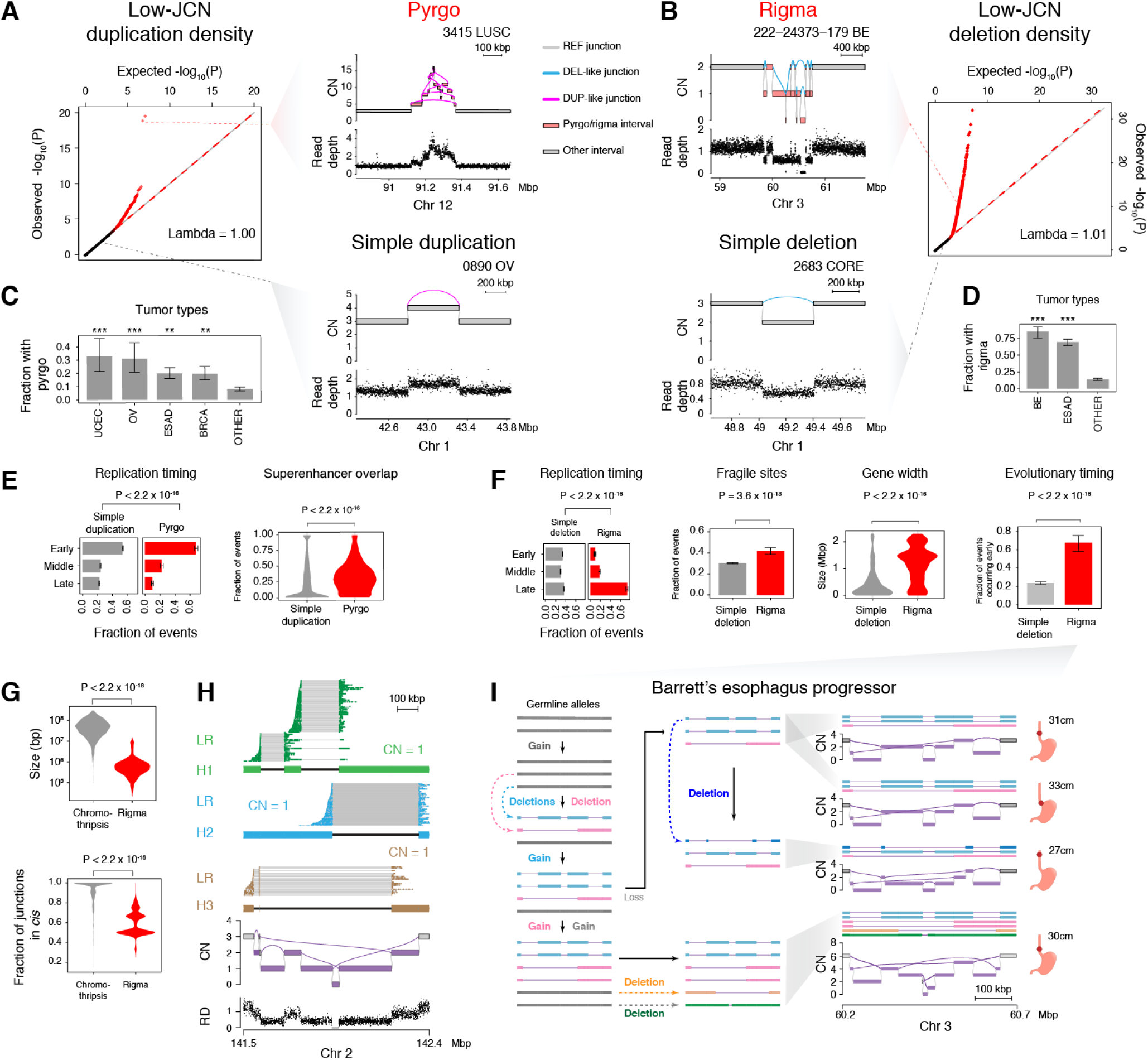
Rigma and pyrgo represent novel patterns of clustered low copy rearrangements. A) Quantile-quantile (Q-Q) plot of observed versus expected p-values illustrating the probability that the observed densities of low copy duplications in 1 Mbp sliding windows falls within the expected gamma-Poisson distribution and expected p-values from a uniform distribution. Red dots indicate sample specific windows that contain density outliers. Top right, an example of a window that contains a high density of DUP-like rearrangements within a sample. We termed this event “pyrgo”. Bottom right, a non-outlier window containing a single DUP-like junction. (B) Right, Q-Q plot similar to (A) instead plotting the observed vs. expected p-values of DEL-like junction density within sliding windows. Top left, an example of a window containing a high density of DEL-like junctions within a sample, an event type we term “rigma”. Bottom left, a non-outlier window containing a DEL-like event. (C) Fraction of samples within tumor types that harbor pyrgo events. Significantly enriched tumor types (compared to all others) marked by asterisks. Significance levels: *** (*FDR* < 1 × 10^*−*3^), ** (*FDR* < 0.01), * (*FDR* < 0.10) (D) Fraction of samples within tumor types that harbor rigma events. See (C) for denoted significance levels. (E) Left, comparison of the fraction of pyrgo footprints vs simple duplication footprints that fall within early, middle and late replicating regions. Right, fraction of pyrgo footprints that overlap with an annotated superenhancer region. (F) Replication timing, comparison of the fraction of rigma footprints and simple deletion footprints that fall within early, middle, and late replicating regions. Fragile sites, comparison of the fraction of rigma with the fraction of simple deletion footprints that overlap with known fragile sites. Gene width, comparison of widths of genes overlapping rigma and deletions. Evolutionary timing, a comparison within the Barrett’s cohort, for which multiple biopsies exist, of the fraction of events that occur early (i.e., in multiple samples from the same patient) in simple deletions and rigma. (G) Top, comparison of the total genomic territory covered by chromothripsis events and rigma events. Bottom, the fraction of rearrangements that occur in *cis* (i.e. on the same predicted haplotype) when the longest possible contigs are inferred from the JaBbA graph. (H) Linked-read sequencing confirming the WGS inferred reconstruction of a rigma event’s junctions occur not in *cis* but on separate haplotypes (i.e. in *trans*). (I) Reconstruction of haplotypes from multiple samples from a single case in the Barrett’s esophagus WGS dataset. P-values obtained by Wald test from ordinal logistic regression (C) or logistic regression (D-F). For (G) p-values were obtained by Wilcoxon rank sum test. Significance thresholded by Bonferroni corrected p-values < 0.05.

We found a significantly increased burden of pyrgo in endometrial, ovarian, breast, and esophageal adenocarcinoma (ESAD), while rigma were enriched in Barrett’s esophagus (BE) cases and ESAD (FDR < 0.1, **Fig. 2C-D**). Compared to simple duplications, pyrgo accumulated in early replicating regions (*P* < 2.2 × 10^−16^, *OR* = 1.36, z-test, ordered logistic regression) and superenhancers as defined in (Hnisz et al., 2013) (*P* < 2.2 × 10^−16^ *OR* = 8.09, z-test, logistic regression) (**Fig. 2E**). In contrast, rigma events were significantly enriched in late replicating regions (*P* < 2.2 × 10^−16^, *OR* = 1.5, ordered logistic regression), fragile sites (*P* = 3.6 × 10^−13^, *OR* = 1.95, z-test, logistic regression), and long genes (*P* < 2.2 × 10^−16^, *OR* = 1.31), relative to simple deletions (**Fig. 2F**). These results show genomic distributions of rigma and pyrgo that are distinct from simple deletions and duplications, respectively. Overall, the results indicate that rigma and pyrgo arise from mutational processes that are distinct from those driving the accumulation of simple deletions and duplications.

### Recurrent hotspots of rigma and pyrgo

To nominate loci that are recurrently targeted by pyrgo and rigma across independent patients, we employed fishHook (Imielinski et al., 2017) while correcting for genomic covariates defined above (replication timing, gene width, superenhancer status) (**Fig. S2B-C**). Applying fishHook to find recurrent pyrgo hotspots across 8,642 unique, previously annotated superenhancers (Hnisz et al., 2013) (see Methods), we found 16 loci were significantly mutated above background (FDR < 0.1) with excellent model fitting (*λ* = 1.03, **Fig. S2B**). This included a superenhancer associated with the oncogene *MYC*, targeted by pyrgo in 14 cases spanning 6 cancer types, including 7 esophageal adenocarcinomas (ESAD). Additional recurrent targets of pyrgo included manually annotated Sanger cancer gene census (CGC) genes associated with GISTIC amplification peaks (**Fig. S2B**).

Applying fishHook to analyze rigma recurrence across 18,794 unique genes, we found 17 genes were significantly mutated above background (FDR < 0.1, *λ* = 1.03) even after correcting for replication timing, gene width, and fragile site status (**Fig. S2C**). Among the top loci surviving false discovery correction, *FHIT, WWOX*, and *MACROD2* represent previously identified hotspots of recurrent CNA loss whose significance has been attributed to genomic fragile sites (Zack et al., 2013; Bignell et al., 2010; Beroukhim et al., 2010; Iliopoulos et al., 2006; Fungtammasan et al., 2012; Cheng et al., 2017). Interestingly, we found numerous CGC genes associated with GISTIC deletion peaks among the rigma targets, including the tumor suppressor *CDKN2A* (**Fig. S2C**). Among the top fishHook hits, *FHIT* is a 2.3 Mbp gene previously nominated as an esophageal adenocarcinoma tumor suppressor which accumulates rigma in 77% of BE and 38% of ESAD cases, as well as other gastrointestinal tumors (**Fig. S2D**,**F**). Taken together, these results suggest that either positive somatic selection or additional genomic features (e.g. chromatin states particular to the cell-of-origin of esophageal cancer) may be driving the recurrence of pyrgo (e.g. *MYC*) and rigma (e.g. *FHIT*) hotspots.

### Rigma gradually accumulate deletions in *trans*

Like rigma, chromothripsis is a clustered rearrangement pattern associated with DNA loss. Comparing the footprints of rigma and chromothripsis events across our set of genome graphs, we found that chromothripsis events were one to two orders of magnitude larger in size (chromothripsis median size: 45.07 Mbp; rigma median size: 0.53 Mbp) (**Fig. 2G, upper panel**, *P* < 2.2 × 10^−16^, Wilcoxon test). In addition, we analyzed the allelic structure that is latent in the JaBbA subgraphs corresponding to chromothripsis and rigma events. Specifically, we searched the subgraph associated with each event for a single path or allele that held the most ALT junctions in *cis* (see Methods). We found that in chromothripsis we were often able to find alleles that placed a higher proportion of the junctions in *cis* relative to rigma (**Fig. 2G, bottom panel**, *P* < 2.2 × 10^−16^, Wilcoxon test). These results were consistent with allelic structure that placed rigma-associated deletion junctions in *trans*. To validate this pattern, we generated 10X Chromium WGS for a rigma-harboring Cancer Cell Line Encyclopedia (CCLE) cell line in our cohort (NCI-H838) to deconvolve alleles in the JaBbA genome graph. Indeed, our allelic reconstruction found evidence for independent linear alleles in *trans* orientation (**Fig. 2H**, see Methods). Each allele was inferred to have a copy number of one, with the superimposed alleles accounting for every copy of every interval and junction in the short read-derived WGS JaBbA subgraph. Of the four DEL-like junctions associated with this rigma, all but one pair was in *trans*.

Given the enrichment of rigma in BE, we probed whether these events occurred early or late in BE evolution. Analyzing 347 multi-regionally, and longitudinally sampled paired biopsies (340 esophageal and 7 gastric) taken from 80 BE cases, we found that a rigma event was significantly more likely than a simple deletion to be found in two or more biopsies (*P* < 2.2 × 10^−16^, *OR* = 6.69, Fisher test) rather than be private to a single biopsy from each patient (**Fig. 2F, right-most panel**). We concluded that rigma are an early feature of Barrett’s esophagus, and may implicate it as an early event in progression to esophageal adenocarcinoma tumorigenesis. Reconstructing the allelic evolution in an early rigma case, we found evidence for successive accumulation of DEL-like junctions, with DEL-like alleles appearing on alleles that had already suffered a previous deletion (**Fig. 2I**). These results suggest that rigma represent a gradual SV mutational process that targets late replicating fragile sites and represents an early event in esophageal adenocarcinoma tumorigenesis.

### Subgraphs of high-JCN junctions reveal genomic typhoons

We then sought to investigate the rearrangement patterns associated with high-JCN junctions (JCN > 7) in our genome graphs. A junction at such an extreme of JCN may evolve through a double minute (DM), breakage fusion bridge cycle (BFBC), or as yet undescribed mechanisms for duplicating already rearranged DNA. To characterize independent amplification events associated with these high-JCN junctions, we first identified 12,327 subgraphs among the 2,798 genome graphs harboring an interval CN of at least twice ploidy (**Fig. 3A**), identifying among these high-level amplicons (amplified clusters within a genome) those that harbor at least one junction with JCN > 7. Among these 1,675 high-level amplicons, we annotated them according to several features: 1) the maximum interval CN in the sub-graph (MICN), 2) the maximum JCN normalized by MICN (MJCN), 3) the summed JCN associated with fold back inversion junctions (INV-like junctions that terminate and begin at nearly the same location in the genome, see Methods) normalized by MICN (FBIJCN), and 4) the total number of high-JCN junctions (NHIGH) in the cluster (**Fig. 3B**).

**Fig. 3.**
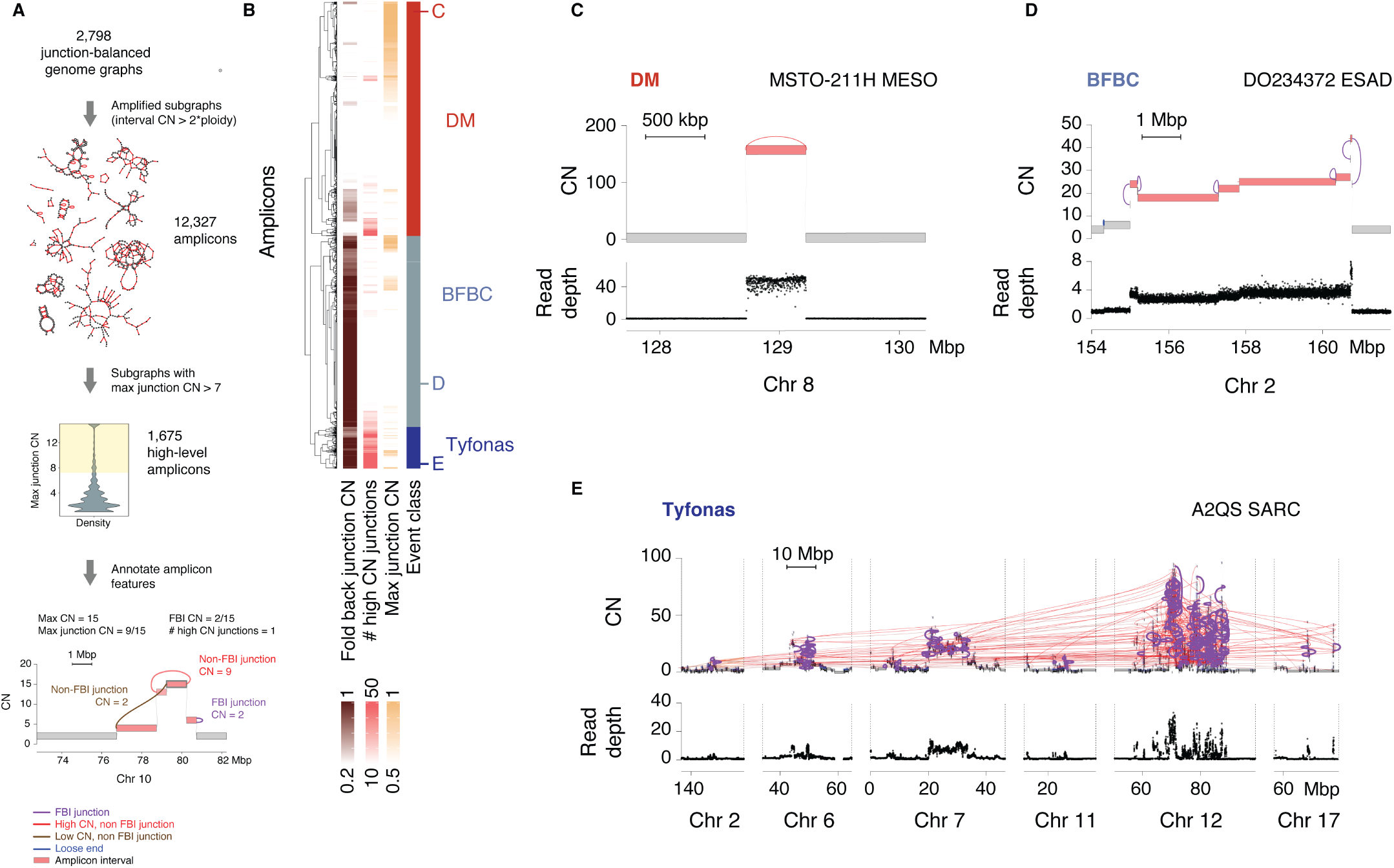
Analysis of amplified subgraphs/amplicons identifies tyfonas. (A) Framework to identify features of complex amplified loci. (B) Heatmap illustrates the separation of three groups of amplicons by hierarchical clustering upon these features. We identified these groups as distinct amplification events: double minutes (DM), breakage fusion bridge cycles (BFBC) and tyfonas. (C) An example of an amplicon we describe as a double minute (DM) event. Top track is the JaBbA-estimated copy number. Bottom, normalized read depth data. (D) Example of an amplicon illustrating typical features of a BFBC and fall within our BFBC grouping. (E) Representative example of an amplicon from the tyfonas group, which is composed of a large number of fold back inversions connected by other SVs through multiple windows.

Hierarchical clustering of amplicons on the basis of these three features yielded three major groups, one associated with low FBIJCN and two associated with high FBIJCN (**Fig. 3B**). We trained a decision tree (see Methods) using the hierarchical cluster labels to derive the feature cutoffs that distinguished these three groups from one another. The low FBIJCN group, contained a subgroup with MJCN near 1, which upon inspection contained amplicons comprising a single high-JCN junction forming a high copy circular path in the graph (**Fig. 3C**), as well as more complex cyclic patterns spanning multiple discontiguous loci. These patterns were most consistent with DM. Among the two high FBIJCN group (FBIJCN > 0.61), there was a class of amplicons with low NHIGH values (< 27), which upon manual review comprised loci with patterns consistent with breakage-fusion-bridge cycles (BFBC), i.e. multiple high copy FBI junction associated with “stairstep” patterns of copy gains (**Fig. 3D**) (Garsed et al., 2014). The third group, contained both high FBIJCN (≥ 0.61) and a large NHIGH count (≥ 27) which comprised dense webs of high-JCN junctions across subgraphs comprising > 100 Mbp of genomic material and often reaching copy numbers higher than 50. We dubbed these extremely large amplicons, which did not fit in a previously defined category, *tyfonas* (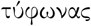, Greek meaning typhoon) (**Fig. 3E**).

### Tyfonas are distinct amplification events from BFBC and DM

Comparing additional features of these high-level amplicon patterns, we found that tyfonas were associated with significantly larger genomic mass (summed interval width weighted by copy number) than either BFBC or DM (tyfonas vs. BFBC: *P* < 2.2 × 10^−16^; tyfonas vs. DM: *P* < 2.2 × 10^−16^, Wilcoxon rank-sum test) and also had a greater MICN (tyfonas vs. BFBC: *P* < 2.2 × 10^−16^; tyfonas vs. DM: *P* < 2.2 × 10^−16^. **Fig. 4A**). While DMs were enriched in glioblastoma, breast cancer, and small cell lung cancer, BFBCs were enriched in lung squamous cell cancer and head neck squamous (FDR < 0.1, Fisher test, **Fig. 4B**). Tyfonas were enriched in sarcoma, breast cancer and melanoma (FDR < 0.1, Fisher test). In particular tyfonas were found in over 80% of dedifferentiated liposarcomas and 40% of acral melanomas, and rarely observed (< 2%) in fibrosarcomas and cutaneous melanomas. All three event types were enriched in ESAD. We analyzed the distribution of BFBC, DM, and tyfonas events across GISTIC peaks containing CGC cancer genes (**Fig. 4C**). DM were most frequently implicated in the amplification of *EGFR* and *NFE2L2*. BFBC were most frequently implicated in *ERBB2, CDK6*, and *CCND1* amplification. Finally, *MDM2, CDK4*, and *NSD1* were the most frequent oncogenic target of *tyfonas*.

**Fig. 4.**
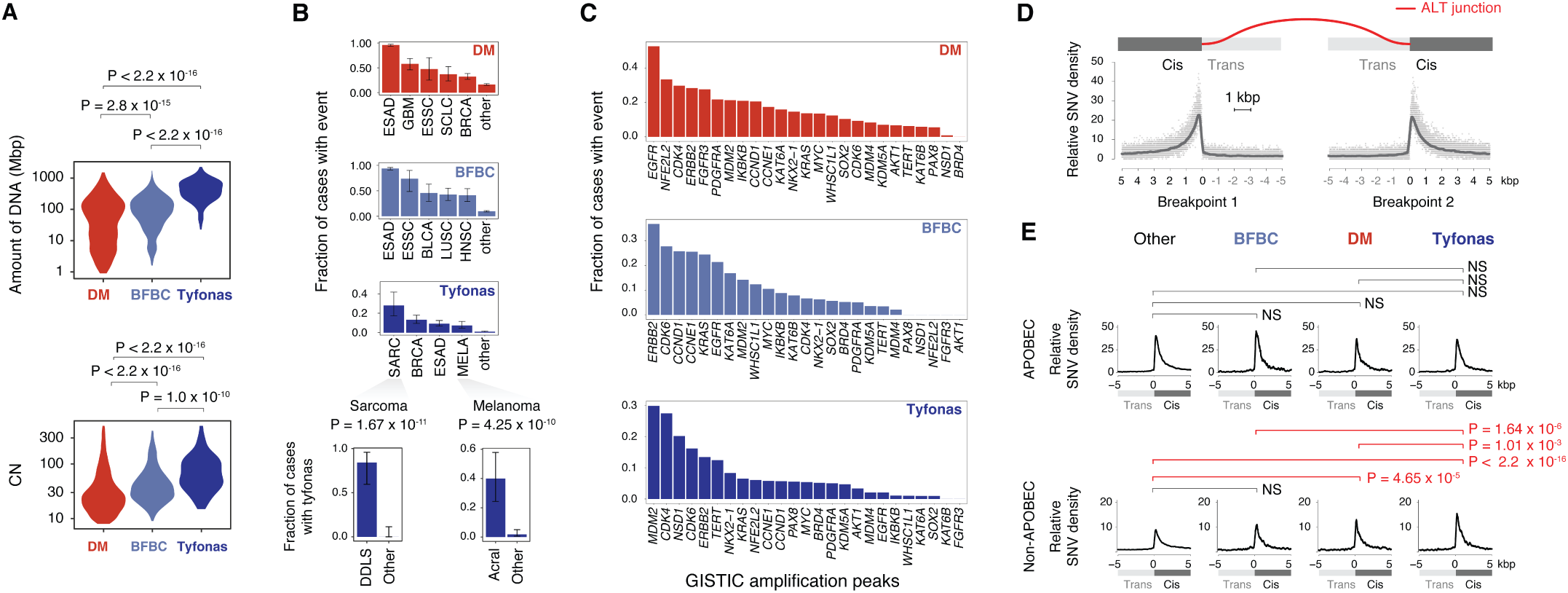
Tyfonas are distinct from DM and BFBC and enriched in non-APOBEC kataegis. (A) Top, distribution of the widths of all intervals covered by each of the three event types weighted by the base-level copy number within each interval. Bottom, the distribution of the maximum interval copy number (MICN), the highest copy number occupied by a unique event interval. (B) Comparison of fraction of cases with each event type amongst tumor types. Particular enrichments of tumor subtypes within sarcomas and melanomas illustrated below (DDLS: Dedifferentiated liposarcoma). (C) Fraction of cases that have overlapping amplification events with driver amplification genes from (Zack et al., 2013), grouped by event type. (D) Breakpoint-centric coordinate system to analyze kataegis near rearrangement breakpoints. On this new axis, every breakpoint is collapsed to the origin (coordinate 0 on x-axis). Top, the *cis* (+ coordinates) sides of the SV have undergone fusion through the rearrangement event (red-colored line), while the *trans* (− coordinates) sides are disconnected from the derivative allele. Bottom, relative SNV density is the count of SNV at every base pair on this axis normalized to the average SNV count from 0 kbp to −5 kbp on this axis. The new axes are shown splitting each rearrangement into each breakpoint side arbitrarily in the dot histograms to illustrate the *cis* and *trans* convention. (E) Relative SNV density on breakpoint-centric coordinates for each of the event types. Top, APOBEC attributed SNV density near breakpoints. Bottom, non-APOBEC attributed SNV density near breakpoints. P-values obtained from Wald test by gamma-Poisson regression comparing *cis* SNV density to *trans* SNV density. Significance determined by Bonferroni correction at a threshold of < 0.05.

### Tyfonas junctions are enriched in non-APOBEC kataegis

To explore whether distinct mutational processes were implicated in the genesis of BFBC, DM, and tyfonas, we examined the patterns of somatic hypermutation around rearrangement breakpoints, also called *kataegis*. Junction breakpoints can be associated with a *cis* side that is fused following rearrangement and a *trans* side that (in the majority of junctions) is lost following rearrangement. Plotting SNV density across all junctions in our dataset with respect to a standard frame (shown in **Fig. 4D**) and normalizing to the density on the *trans* side shows a distinct pattern of hypermutation on the *cis* side which is most prominent in the first 1 kbp. Applying a gamma-Poisson regression model to quantify the enrichment of mutation counts in the first *cis* 1 kbp relative to the first 5 kbp territory on the *trans* side (from 0 kbp to 5 kbp away from the breakpoint) allows us to statistically assess the presence of kataegis and differences in the degree of kataegis between junction and SNV categories (see Methods).

Kataegis has been classically associated with APOBEC mutagenesis, which can be defined using previously defined COSMIC Signatures 2 and 13 (Alexandrov et al., 2013; Nik-Zainal et al., 2012). Strikingly, our results demonstrate statistically significant kataegis both within and outside of the APOBEC context (**Fig. S3A-D**). Analyzing APOBEC-associated SNV signatures, we found no significant differences in kataegis between BFBC, DM, and tyfonas, and baseline junctions (i.e. those not associated with a high-level amplicons) (**Fig. 4E, top row**). However, we found that DM (*P* < 2.2 × 10^−16^, *RR* = 1.27, z test, gamma-Poisson regression) and tyfonas (*P* < 2.2 × 10^−16^, *RR* = 1.62, z test, gamma-Poisson regression) were enriched in non-APOBEC kataegis relative to baseline, with tyfonas showing a statistically significant increase in kataegis relative to DM (*P* = 1.01 × 10^−3^, *RR* = 1.28) and BFBC (*P* = 1.64 × 10^−6^, *RR* = 1.43) (**Fig. 4E, bottom row**). These results show that tyfonas are enriched in a previously undescribed mutational process causing non-APOBEC hypermutation around junctions.

### Genome graph features define distinct tumor clusters

Tallying counts of genome graph-derived SV patterns across previously characterized simple (deletions, duplications, inversions, inverted duplications, translocations) and complex (chromothripsis, chromoplexy, templated insertion chains) event classes as well as the novel patterns introduced above (pyrgo, rigma, tyfonas) allowed us to group 2,798 tumor/normal pairs into 13 distinct clusters (**Fig. 5A, Fig. S4A**). We labeled these clusters according to the particular event types that were enriched among them (except for the Quiet cluster which displayed a dearth of events), including clusters associated with previously identified event types such as the CT (chromothripsis) and CP (chromoplexy) cluster. Consistent with previous reports, the CT cluster was significantly enriched in prostate adenocarcinoma (PRAD) (*P* = 6.16 × 10^−10^, *OR* = 4.05, z-test, logistic regression) and glioblastoma multiforme (*P* = 1 × 10^−4^, *OR* = 3.00, z-test, logistic regression) (**Fig. 5B**). Similarly, we found that the CP (chromoplexy) cluster was significantly enriched in PRAD (*P* = 3.21 × 10^−10^, *OR* = 4.85). DDT tumors (defined by deletions, duplications, and templated insertion chains) were enriched in breast and ovarian cancer as well loss of function mutations in several genes notably involved in DNA repair (**Fig. 5C**): More than 30% of cases in the DDT cluster harbored loss of function lesions in *BRCA1*, which was statistically significantly enriched above baseline even after correcting for tumor type (*P* = 3.20 × 10^−7^, *OR* = 165.67). In addition, we found that DDT tumors were also enriched in somatic loss of function mutations in *RB1*, even after correcting for tumor type (*P* = 1.95 × 10^−5^, *OR* = 219.20).

**Fig. 5.**
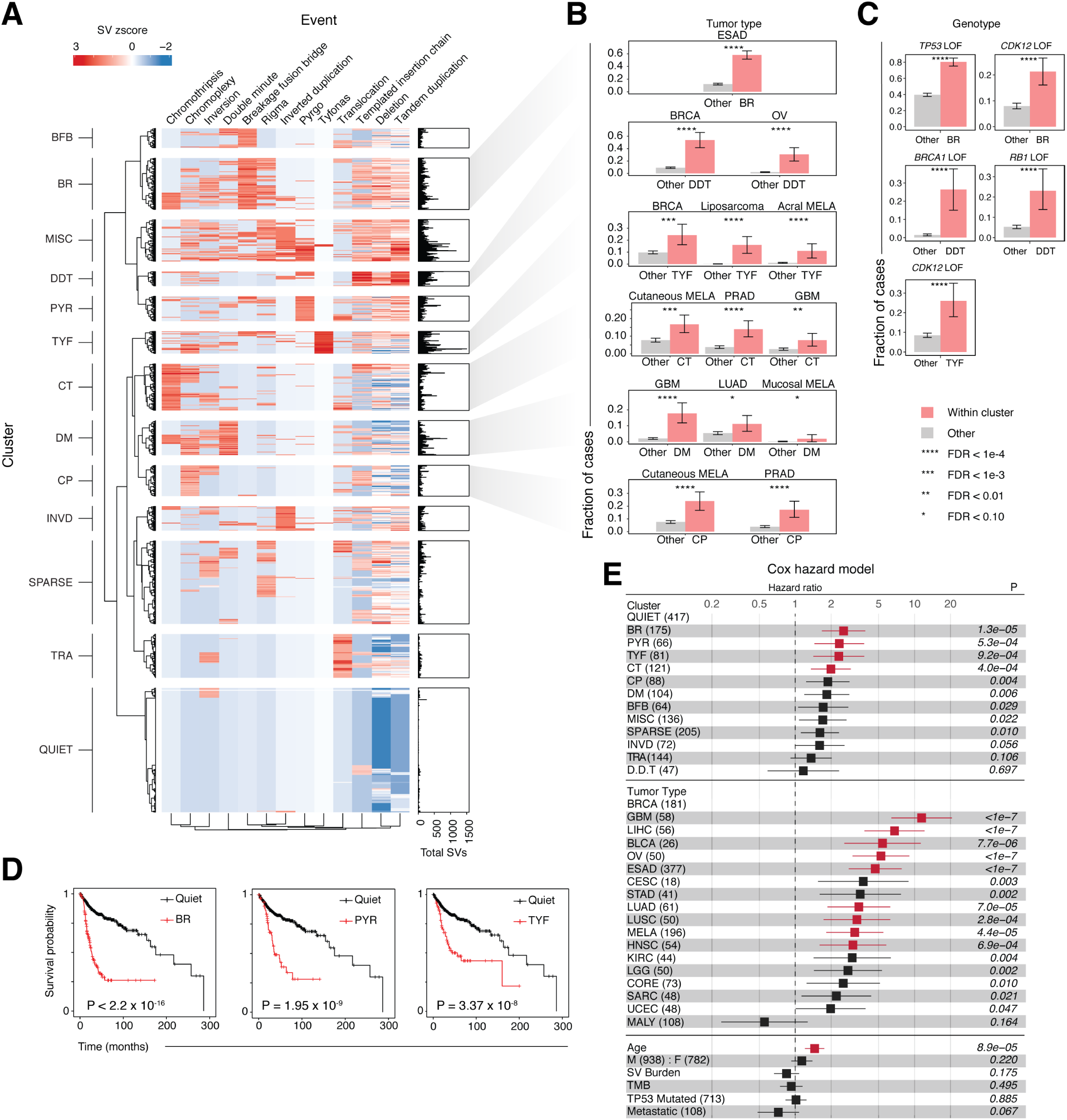
Genome graph-derived SV features define biologically distinct and prognostically important clusters. (A) 13 distinct clusters of cases illustrated by a heatmap of negative binomial-distributed Z-scores of junctions attributed to each event type detected from JaBbA graphs across 2,517 unique patients. Abbreviations indicate cluster names which identify the enriched event within the cluster (BFB, breakage fusion bridge; BR, BFB and rigma; MISC, miscellaneous; DDT, deletion, duplication, and TIC; PYR, pyrgo; TYF, tyfonas; CT, chromothripsis; DM, double minute; CP, chromoplexy; INVD, inverted duplications; TRA, translocations. (B) Fraction of cases in tumor types significantly enriched in selected clusters. Red bars indicate fraction of cases within a cluster for the labeled tumor type. Grey bars indicate fraction of cases outside of a given cluster for the labeled tumor type. (C) Fraction of cases with loss of function in select genes within select clusters. Red bars indicate fraction of cases within a cluster that harbor loss of function for the given gene. Grey bars indicate fraction of cases outside of the cluster that have loss of function for a given gene. Significance in (B) and (C) determined by Wald test from Bayesian logistic regression models, thresholded at *FDR* < 0.10. Significance levels: **** (*FDR* < 1 × 10^*−*4^), *** (*FDR* < 1 × 10^*−*3^), ** (*FDR* < 0.01), * (*FDR* < 0.10) (D) Kaplan Meier curves showing reduced survival for those cases that fall within BR, PYR and TYF clusters (i.e. clusters dominated by BFBC/rigma, pyrgo, and tyfonas events, respectively). P-values obtained via log-rank test. (E) Cox hazard ratios comparing the relative risk cases within each cluster to the QUIET cluster comprised of cases harboring few rearrangement events. Cox regression results shown for covariates known to associate with survival: age, tumor type (compared to BRCA as the reference tumor type), sex, metastasis, tumor mutational burden, overall SV burden, and *TP53* loss of function status. P-values shown are not corrected. Significantly associated variables (Bonferroni *p* < 0.05) colored in red. Bonferroni correction performed within each variable group separately (i.e. within Cluster, and within Tumor Type separately).

Inspection of the clustered heatmap in **Fig. 5A** showed that the novel event categories introduced in this study (pyrgo, rigma, tyfonas) were distributed independently relative to previously identified complex event types (DMs, BFBCs, Chromothripsis, Chromoplexy) Supporting this assertion, we found three clusters (BR, PYR, TYF) defined by the enrichment of at least one of these novel event types. The BR (BFBC, Rigma) cluster was almost 60% composed of ESAD cases (*P* < 2.2 × 10^−16^, *OR* = 9.87, z-test, Bayes logistic regression) (**Fig. 5B**). BR-cluster tumors were also significantly enriched in somatic *TP53* mutations (*P* < 2.36 × 10^−9^, *OR* = 6.26) (**Fig. 5C**) as well as germline *TP53* mutations (*P* = 1.4 × 10^−4^, *OR* = 5.51) (**Fig. S4B**). In addition, tumors in the BR cluster were significantly enriched in *CDK12* loss of function mutations (*P* = 1.58 × 10^−5^, *OR* = 3.39). The TYF (tyfonas) cluster was enriched in breast cancer, dedifferentiated liposarcoma, and acral melanoma. In contrast, cutaneous melanomas were enriched in the CT cluster, while mucosal melanomas were enriched in the DM cluster. The TYF cluster also harbored an increase in *CDK12* loss of function mutations relative to baseline (**Fig. 5C**).

We then asked whether the observed clusters demonstrated significant overall survival differences. Indeed, Kaplan-Meier analysis showed that several clusters were significantly associated with poor survival relative to the Quiet cluster, including BR, PYR, and TYF (**Fig. 5D**, FDR < 0.1, log rank test). This negative prognostic impact was still statistically significant for BR (*P* = 1.46 × 10^−4^, *HR* = 2.54, likelihood ratio test, Cox regression), PYR (*P* = 6.08 × 10^−3^, *HR* = 2.34), and TYF (*P* = 9.70 × 10^−3^, *HR* = 2.32) even after adjusting for the distribution of tumor types, overall SV burden, tumor mutation burden (TMB), *TP53* mutation status, and metastatic vs. primary sample status in a multivariate Cox regression analysis (**Fig. 5E**). These results show that the novel genome-graph-derived features introduced in this study define biologically distinct tumor clusters which are enriched in specific DNA repair defects and show distinctly poorer prognosis, even after taking into account known determinants of survival.

## Discussion

Our genome graph based framework establishes the topology of JCN as an important signal for classifying complex SV patterns in cancer. By leveraging statistical deviations in the genomic distribution of JCN and the topology of junction-balanced genome graphs, we nominate three novel event types: pyrgo, rigma, and tyfonas. The regional, genotypic, and tumor type enrichment of these previously undescribed complex SV patterns is consistent with these being the product of specific mutational processes, distinct from those driving previously identified complex rearrangement patterns (chromothripsis, chromoplexy, BFBC). These may either be driven by cell-of-origin features (e.g. chromatin state, replication timing), tissue type specific mutagens, or the inactivation of specific genome integrity pathways. We provide the full set of 2,798 annotated genome graphs as a data portal http://mskilab.com/gGraph which can be explored through our custom gGnome.js genome graph browser.

Our data show that rigma likely arise from an early and ongoing accumulation of deletions at large and late replicating genes. These locations are enriched, but not perfectly associated, with previously annotated fragile sites defined through cell culture experiments involving exposure to aphidicolin, cytosine analogs, and dNTP depleting compounds (Schwartz et al., 2006) and mapped by cytogenetics (Fungtammasan et al., 2012) or exon trapping (Ohta et al., 1996). An intriguing possibility is that a subset of these rigma represent additional previously unannotated genomic fragile sites, which may be important for the study of other diseases (e.g. development delay or autism). The recurrence of rigma at *FHIT* and other fragile-site associated genes (e.g. *WWOX, MACROD2*) suggests that replication timing and gene size, although highly correlated with chromosomal fragility (Mrasek et al., 2010; Iliopoulos et al., 2006) do not fully account for their accumulation in hotspots. Though some of these rigma hotspots may be *bona fide* drivers, another possibility is that there are uncharacterized or cell-type specific features of the cell-of-origin chromatin state that causes these events to recur at particular genes. The preference of rigma for ESAD (and more broadly GI cancers) suggests that these unique chromatin features may be found through the analysis of cell types in the healthy GI epithelium (e.g. through single cell approaches, or cell sorting and chromatin profiling). These findings complement recent discoveries demonstrating unprecedented somatic genomic complexity in normal epithelial tissues (Lee-Six et al., 2019; Martincorena et al., 2018; Brunner et al., 2019).

Our analysis of high-level amplicons identifies three distinct groups of copy-amplifying events, which we label broadly as BFBC, DM, and Tyfonas. A key question is whether tyfonas are extrachromosomal like DM (Verhaak et al., 2019), or integrated into chromosomes like BFBC. We found that both the maximum interval CN of tyfonas and the genomic footprint of these events is an order of magnitude higher than DM or BFBC. The enrichment of tyfonas in over 80% of dedifferentiated liposarcomas and clustering to the *MDM2* / *CDK4* locus on chromosome 12q suggests that these events represent supernumerary ring chromosomes described in classic cytogenetics studies of this disease (Reimann and Fletcher, 2008). A previous cell line study of dedifferentiated liposarcomas suggested that these characteristic amplifications may arise as extrachromosomal DNA which grows to a large size and accumulates additional junctions through a “circular BFBC” mechanism, before eventually acquiring a centromere and becoming a neochromosome (Garsed et al., 2014). Interestingly, supernumerary rings have not been previously associated with acral melanoma, though cytogenetics studies have been limited in this disease.

Our discovery of non-APOBEC driven hypermutation around DNA rearrangement breakpoints has not been (to our knowledge) previously reported. Though our analyses employ one of the several definitions of APOBEC driven mutagenesis (COSMIC Signatures 2 and 13), we obtain very similar findings with broader or alternate definitions of APOBEC mutagenesis (e.g. GC-strand coordinated clusters (Roberts et al., 2013), TpC mutations (Nik-Zainal et al., 2012)) (**Fig. S3A**). Intriguingly, if tyfonas and DM are both extrachromosomal, the enrichment of non-APOBEC kataegis in these events (but not BFBC) may represent the footprint of a novel SNV mutational process that affects rearrangement junctions arising in extrachromosomal DNA. The relative enrichment of non-APOBEC kataegis in tyfonas relative to DM may then reflect the higher burden of (late) extrachromosomal-derived junctions in these events.

The enrichment of tyfonas in acral but not cutaneous melanomas provides intriguing context for the observation that both of these melanoma subtypes are responsive to immune checkpoint inhibition (ICI) therapy (Shoushtari et al., 2016). While the ICI responsiveness of cutaneous melanoma is attributed to the accumulation neoantigens arising from UV-driven SNVs, acral melanomas harbor few SNVs because they arise in sun-protected regions (e.g. feet). An intriguing possibility is that the massive degree of rearrangement and amplification induced by tyfonas events may serve to generate neoantigens and thus explain the responsiveness of acral melanomas to ICI. If so, the analysis of genome graphs and identification of tyfonas could be applied clinically in other tyfonas-harboring tumor types (e.g. small cell lung cancer) as a WGS biomarker to nominate patients for ICI therapy.

Our genome graph-based cancer classification provides novel links between specific genome integrity pathways and SV evolution. The significant enrichment of *BRCA1* mutations in the DDT cluster suggests a previously unidentified link between homologous repair deficiency and templated insertion chains. Previous mouse model and cell line studies have linked BFBC evolution to *TP53* loss (Bianchi et al., 2019; Gisselsson et al., 2000). The enrichment of both inherited and acquired *TP53* mutations in the poor-prognosis and esophageal adenocarcinoma-associated BR cluster provides some of the first evidence in human disease linking *TP53* to the evolution of BFBC. Interestingly, the inclusion of BR cluster membership corrects for the negative prognostic impact of *TP53* mutation status in our multivariate model of survival. These results may indicate that *TP53* mutations may drive the evolution of a particularly aggressive subtype of esophageal adenocarcinomas, which is marked by the accumulation of rigma and BFBC.

Our study demonstrates the importance of JCN and graph topology in the characterization of complex SV in cancer. However, a considerable fraction of the junctions in our dataset remained unclassified with respect to any of the 14 complex rearrangement event patterns that we have catalogued and/or introduced (40%, **Fig. S5A**). Some of this gap can be attributed to missing junctions in short read WGS, which introduce loose ends into the graph and fracture the subgraph structure around copy number alterations. Missing junctions occur due to sampling (ie inadequate purity and/or read depth) or mappability limitations in the sequencing platform (junctions arising in repetitive sequence) and can be overcome with additional sequencing as well as longer molecules (Sedlazeck et al., 2018).

Our genome graphs provide a starting point for rigorous classification of complex SV patterns, but only lend partial insight into mechanism. For example, it is unclear whether pyrgo represent another facet of the “tandem duplicator phenotype” (Menghi et al., 2016; Viswanathan et al., 2018; Willis et al., 2017) or arise from a completely independent mechanism. Additional mechanistic insight into the genesis of complex SV will likely require the consideration of junction *phase*, as shown with our 10X Chromium WGS analysis of rigma. Though our JaBbA-derived genome graphs provide a starting point to deconvolve phased alleles across complex loci in short read WGS, these locus reconstructions usually yield multiple solutions in the absence of long-range genome profiling data (10X Chromium WGS, Pacific Biosciences, Oxford Nanopore Technologies, Hi-C, BioNano) (Sedlazeck et al., 2018). In addition, integration of SV (and/or SNV) derived features using topic models or non-negative matrix factorization may help link linear combinations (i.e. signatures) of complex SV types to environmental exposures and DNA repair defects (Macintyre et al., 2018; Funnell et al., 2019; Alexandrov et al., 2013; Nik-Zainal et al., 2016). Comprehensive characterization of the long-range allelic phase of complex SV across large clinically annotated cohorts, leveraging multi-regional and/or single cell sequencing, will be essential to gain insight into the mutational mechanisms underlying SV evolution.

### Software Availability

Software used in this paper can be found in the following GitHub repositories:

- https://github.com/mskilab/JaBbA
- https://github.com/mskilab/gGnome
- https://github.com/mskilab/gGnome.js

## ACKNOWLEDGEMENTS

M.I. thanks Dan Landau for critical and insightful discussions. M.I. thanks Matthew Meyerson and Cheng-Zhong Zhang for helpful feedback early in the development of JaBbA. M.I. thanks Ron Zeira, Ben Raphael, and Tomasz Imielinski for feedback on the JaBbA mathematical formulation. The authors thank Christopher Black and the NYGC high performance compute team for their support during this project. M.I., X.Y, J.M.B., A.D., J.R., H.T., E.A., and Z.G. are supported by M.I.’s Burroughs Wellcome Fund Career Award for Medical Scientists, Doris Duke Clinical Foundation Clinical Scientist Development Award, Starr Cancer Consortium Award, Melanoma Research Alliance Team Science Award, National Institutes of Health U24-CA15020, and Weill Cornell Medicine Department of Pathology and Laboratory Medicine startup funds. K.H. is supported by a NIH/NCI F31 Graduate Research Fellowship (F31-CA232465). M.D. is supported by an NIH F32 training grant given to the Tri-Institutional Computational Biology and Medicine PhD Program. R.B. received funding from the Fund for Innovation in Cancer Informatics. B.J.R., T.P., X.L., P.G., C.S., K.O., M.K., and L.S. are supported by NIH funding (P01-CA9195). The IBM-NYGC Cancer Alliance project was supported in part by a grant from the IBM corporation (IBM Watson Health) to the New York Genome Center, New York Genome Center philanthropic funds and Rockefeller University grant UL1 TR000043 from the National Center for Advancing Translational Sciences (NCATS), and the National Institutes of Health (NIH) Clinical and Translational Science Award (CTSA) program.

## AUTHOR CONTRIBUTIONS

These contributions follow the Contributor Roles Taxonomy guidelines: https://casrai.org/credit/. Conceptualization: X.Y., K.H., J.B., J.W., R.B, T.D.L., J.M, J.S., Na.R., J.S.R.-F., S.P., M.I.; Data curation: X.Y., K.H., J.B, A.D., J.R., M.D, H.T., Z.G., K.E., K.O.W., K.A., M.S., A.K.E, V.F., M.O.F., M.G., F.H., J.M., Ni.R., K.M.O, C.A.S., M.K.K, L.P.S., P.C.G., T.G.P.à, X.L., D.W., A.S., J.M.M, M.I.; Formal analysis: J.B., K.H., X.Y, A.D., M.I.; Funding acquisition: R.D.,P.G., B.R., O.E., M.I.; Investigation: X.Y, J.B., K.H., A.D., M.I.; Methodology: X.Y., J.B., K.H., A.D., D.K., E.R., B.M, M.I.; Project administration: M.I.; Resources: O.E., R.D., M.G., F.H., T.D.L, M.Z., N.R., T.P., P.G., R.S., X.L., B.R., M.I.; Software: X.Y., J.B., C.X., M.I.; Supervision: M.I.; Validation: X.Y., J.B, K.H., A.D., M.I.; Visualization: X.Y., J.B., K.H., C.X., M.I.; Writing – original draft: M.I.; Writing – review & editing: all authors.

## Methods

### Pan-cancer data curation

Out of the 2,833 WGS from total tumor/normal pairs and cell lines covering 31 primary tumor types (**Table 1**), 2,274 were obtained from The Cancer Genome Atlas (TCGA), the International Cancer Genome Consortium (ICGC), the Cancer Cell Line Encyclopedia (CCLE), or other previously published data (described below). In total, 3,882 BAMs were downloaded from the respective institutional or publicly available repositories. Criteria for inclusion into this study were as follows: i) BAMs from only non-low pass WGS, ii) BAMs must be aligned to GRCh37/hg19, and iii) both tumor and normal non-low pass WGS must exist per pair except for cell lines. Previously published WGS cohorts included in this study were: 183 ICGC melanoma cases (Hayward et al., 2017), 49 ICGC lung adenocarcinomas (Lee et al., 2019), 122 ICGC breast cancers (Nik-Zainal et al., 2016), and 422 ICGC esoaphgeal adenocarcinomas (Frankell et al., 2019), 55 prostate cancers (Baca et al., 2013), and also 326 unpaired CCLE cell lines (Barretina et al., 2012). Raw sequencing data was obtained either from either public repositories with the proper permissions granted through dbGaP (for TCGA WGS BAMs) or through the relevant Data Access Committees (for ICGC WGS BAMs). TCGA WGS BAMs were downloaded from the Genomic Data Commons (GDC, https://portal.gdc.cancer.gov/legacy-archive). All ICGC BAMs were downloaded from the European Genome-Phenome Archive (EGA, https://ega-archive.org). The additional 49 lung adenocarcinoma cases (Lee et al., 2019) and 55 prostate cancers (Baca et al., 2013) were obtained through data access agreements with the relevant institutions involved.

Standard WGS data from two additional cell lines, G15512.HCC1954 (58X) and its paired normal G15512.HCC1954BL (71X), were also downloaded from the GDC in BAM format, as a part of the TCGA mutation/variation calling benchmarking 4 dataset. The *uuid* for the file retrieval were *6d8044f73f63487c9191adfeed4e74d3* and *34c9ff85c2f845dcb4aafba05748e355*, respectively. For haplotype reconstruction, we also generated 10X linked-read sequencing data for these cell lines (see below for protocol) (10X Genomics, Pleasonton, CA).

All collected standard short-read WGS reads were aligned using the Burrows-Wheeler Aligner (Li and Durbin, 2009) (via `bwa mem` or `bwa aln` settings) to the GRCh37/hg19 reference. Harmonized analytic pipelines (e.g. mutation calling, structural variant calling, and graph genome modeling) were then applied to these data (described in detail below).

### Sample collection

We collected 825 samples, including 546 tumor samples, from 232 patients across three study cohorts: the IBM-NYGC Cancer Alliance (CA), Weill Cornell Englander Institute for Precision Medicine (EIPM), and the Fred Hutchinson Barrett’s Esophagus project (FHBE). Sample characteristics of these cohorts are summarized in **(Table S1)**.

In the CA study, clinically annotated frozen tumor and matched normal (blood, adjacent) samples were collected from 116 consented participants with pathologically verified diagnoses spanning 18 primary tumor types as part of a collaboration spanning nine academic medical institutions in the New York City area, including Memorial Sloan Kettering Cancer Center, New York University, Stony Brook University Hospital, Lenox Hill, Northwell Health, Columbia University, Montefiore, Cornell, and led by the New York Genome Center.. Tumor and normal paired samples were submitted for precision oncology evaluation, including whole exome and whole genome sequencing and interpretation, at the New York Genome Center. This study was approved by a central institutional review board (IRB), Biomedical Research Alliance of New York, and by local IRBs, including Stony Brook University and Northwell Health.

In the EIPM study, tumor and matched normal (blood, normal adjacent) from 83 consented EIPM study participants spanning 12 tumor types were collected as part of a precision WGS oncology pilot study at Weill Cornell Medicine. This study was approved by an Weill Cornell Medicine Institutional Review Board.

In the FHBE study, clinically and pathologically annotated samples from 80 consented BE patients were collected at two time points per patient. At each time point, endoscopic biopsies were taken at two different distances from the esophagogastric junction (GEJ), yielding four samples per patient. Additional biopsies and time points were collected for 10 patients. 62 blood sample and 25 non-pathological gastric tissues were collected as normal controls.

Additional cell lines were obtained for validation and benchmarking studies. Cell lines HCC1143 (cat. no. CRL-2321), HCC1143BL (CRL-2362), HCC1954 (CRL-2338), HCC1954BL (CRL-2339), and NCI-H838 (CRL-5844) (available through ATCC, Manassas, VA) were cultured in ATCC-formulated RPMI-1640 Medium (ATCC 30-2001) with a final concentration of 10% fetal bovine serum (ATCC 30-2020).

### Whole genome sequencing

Library preparation and whole genome sequencing for all sequenced cohorts (CA, EIPM, FHBE) was performed at the New York Genome Center (NYGC) (New York, NY) to a target 80X tumor and 40X normal coverage. For the CA and EIPM studies and HCC1143/HCC1143BL cell lines, short-read genomic DNA library preparation was performed with the KAPA Library preparation kit in accordance with manufacturer’s protocols (Kapa Biosystems, Wilmington, MA). In the FHBE study, genomic DNA was extracted from 427 biopsies (comprising 340 Barrett’s esophagus/normal pairs, and 7 gastric/normal pairs). 415 out of 427 libraries were prepared the TruSeq DNA PCR-free Library Prep Kit (Illumina, San Diego, CA). For the remaining 12 libraries, the TruSeq DNA Nano Library Prep Kit (Illumina) was used. All processing adhered to manufacturer’s protocols. Quality control was assayed for the final libraries with the Agilent 2100 Bio-analyzer by using the DNA 1000 chip (Agilent Technologies, Santa Clara, CA). QC determined that libraries contained an average peak height (fragment size) of ≥400 base pairs. Libraries were sequenced on HiSeq X machines (Illumina) to genrerate paired-end 2×150 base pair reads. Reads were aligned to the GRCh37/hg19 reference using Burrows-Wheeler aligner software (Li and Durbin, 2009) (`bwa aln`, v.0.7.8). Best practices for post-alignment data processing were followed through use of `Picard` (https://broadinstitute.github.io/picard/) tools to mark duplicates, the GATK (v.2.7.4) (https://software.broadinstitute.org/gatk/) `IndelRealigner` module, and GATK base quality recalibration.

### 10X Chromium linked-read whole genome sequencing

Three cell lines, HCC1954, HCC1954BL, and NCI-H838, were subjected to 10X Chromium linked-read whole genome sequencing.

High molecular weight (HMW) genomic DNA (gDNA) was extracted using Qiagen MagAttract HMW DNA Kit (Qiagen, Germany) according to the suggested protocol. Approximately 2 million fresh cells were lysed, HMW gDNA was captured by magnetic particles MagAttract Suspension G, then the magnetic particles with HMW gDNA was washed in wash buffer and eluted in EB Buffer (10 mM Tris-HCl, pH 8.5). The HMW gDNA had a mode length at 50 kb and max length 200 kb, as estimated on a separate 75V pulse-field gel electrophoresis (BluePippin 5-430kbp protocol).

The 10X sequencing library preparation was done using Chromium Genome Library Kit v2 (Lot 152527, 10X Genomics) following the protocol ChromiumTM Genome v2 Protocol Time Planner. 1 ng of extracted HMW gDNA was used to prepare a 150 bp (insert size) paired-end library, with an average fragment length of 625 bp (ranging from 300 to 2000 bp, measured on Bio Analyzer High Sensitivity DNA Kit, Agilent). The prepared library was sequenced on Illumina NovaSeq 6000 Sequencing system with S4 flow cells, to average read depth of about 45X and physical coverage of 167X for HCC1954, 60X and 143X for HCC1954BL, 33X and 173X for NCI-H838. All the 10X linked reads were aligned with Long Ranger (v2.1.3, 10X Genomics).

### JaBbA mathematical formulation

Below, we formally define a reference genome, genome graph, and junction-balanced genome graph (JBGG). We show how we construct genome graphs from junctions and breakpoints obtained through the analysis of cancer whole genome sequences (WGS). We then define the `JaBbA` algorithm to infer JBGGs by fitting integer vertex and edge weights to high-density WGS read-depth data through the solution of a mixed integer quadratic program (MIQP).

### Reference genome

Let the reference genome 𝒞 comprise *c* pairs of strings, labeled *C*^*i*^ and *C*^−*i*^, *i* ∈ 1, …, *c*. Each string pair *i* ∈ 1, …, *c* is called a *chromosome*, and each string in that pair is called a *strand*. We use *C*^*i*^ and *C*^−*i*^ to refer to “positive” and “negative” strands of chromosome *i*, each having length *L*_*i*_ ∈ ℕ. We use brackets to refer to substrings on these strands e.g. *C*^−*i*^[*q, r*] refers to the substring beginning at position *q* and ending at position *r* (inclusive) where *q* ≤ *r* ∈ 1, …, *L*_*i*_. We also use 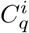 as a shorthand for *C*^*i*^[*q, q*]. Every position 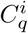 has a “reverse complement” (RC) position 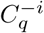. Similarly, every substring *C*^*i*^[*q, r*] has an RC substring *C*^−*i*^[*q, r*].

### Breakpoints and junctions

To describe a rearranged and copy number altered reference genome, we partition 𝒞 according to a collection of *breakpoints* ℬ. We also define a set of *junctions* 𝒜 representing alternative adjacencies between a set of the breakpoints in ℬ. Each *B*^*i*^ ∈ ℬ, *i* ∈ 1, …, *c* is an ordered and unique sequence of integer coordinates 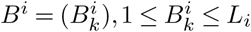 on chromosome *i*, where 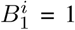 and 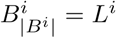. Each junction *A* ∈ 𝒜 is a tuple 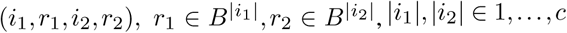 representing a (3’-5’ phosphodiester) bond between the position 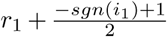 on chromosome / strand 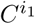 and position 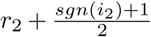 on chromosome / strand 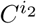. For every adjacency *A* = (*i*_1_, *r*_1_, *i*_2_, *r*_2_) ∈ 𝒜 we require 𝒜 to contain the reverse complement adjacency *Ā* = (−*i*_2_, *r*_2_, −*i*_1_, *r*_1_). The adjacencies in 𝒜 are “alternative” relative to a set of “reference adjacencies” ℛ implied by *B*^*i*^, comprising tuples 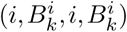 and 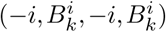 for each breakpoint 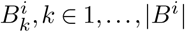 in each chromosome *i* ∈ 1, …, *c*.

### Genome graph

A genome graph is a directed graph *G* = (*V, E, ψ, ϕ*) whose vertices *v* ∈ *V* represent strands of DNA sequences, and whose edges *e* = (*v*_1_, *v*_2_) ∈ *E*(*G*) represent genomic adjacencies (i.e. 3-5’ phosphodiester bonds) joining two DNA sequences, where *v*_1_, *v*_2_ ∈ *V* (*G*) **Fig. S1A**. The mapping *ψ* : *V* (*G*) ∪ *E*(*G*) → {‘*I*’, ‘*L*’, ‘*T* ‘, ‘*R*’, ‘*A*’} is a partition of vertices and edges into key “classes”, which we refer to with subscripts, e.g. *V*_*I*_ (*G*) represents the set of vertices *v* for which *ψ*(*v*) = ‘*I*’. A second mapping *ϕ*(*v*) = (*i, q, r*) maps vertices *v* ∈ *V*_*I*_ in *G* to tuples of integer coordinates representing substrings *C*^*i*^[*q, r*] of reference genome sequence 𝒞. For shorthand, we use *V* = *V* (*G*) and *E* = *E*(*V* (*G*)) when the context is obvious. We also use *E*(*v*) ⊆ *E*(*V*) to refer to the edges associated with a single vertex *v*, and refer to the incoming (“5’”) and outgoing (“3’”) edges of a vertex *v* as *E*^−^(*v*), *E*^+^(*v*) ⊆ *E*(*v*), respectively.

The set *V* is partitioned to “internal” vertices *V*_*I*_ and sequence “ends” *V*_*N*_, where *V* = *V*_*I*_ ∪ *V*_*N*_. The ends in *V*_*N*_ = *V*_*T*_ ∪ *V*_*L*_ further comprise reference ends *V*_*T*_ and “loose” ends *V*_*L*_. The set of edges *E* is partitioned into reference edges *E*_*R*_, alternate edges *E*_*A*_, and loose end edges *E*_*L*_ connecting internal and loose end vertices, where *E* = *E*_*R*_ ∪ *E*_*A*_ ∪ *E*_*L*_. *E*_*R*_ also contains reference edges connecting (telomeric) internal vertices *V*_*I*_ with their reference-adjacent ends *V*_*T*_. We apply subscript notation to refer to incoming and outgoing edges of vertices, e.g. 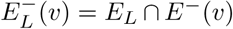 and 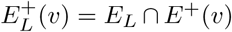 to denote the (single) loose end edge that is upstream and downstream of a vertex *v*, respectively.

### Junction-balanced genome graph

We define a mapping *κ* : {*V*_*I*_ ∪ *E*} → ℕ of non-negative integer copy number (CN) to vertices and edges of *G*, where *κ*(*v*), *v* ∈ *V*_*I*_ and *κ*(*e*), *e* ∈ *E* represent the CN of vertex *v* and edge *e*, respectively. The principle of *junction balance* constrains the CN of every vertex to be equal to the sum of its incoming edges and the sum of its outgoing edges. Formally, the junction balance constraint is stated as follows:

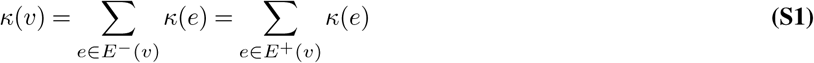

In addition we require the CN *κ* to obey *skew-symmetry*, which means that every vertex must have the same copy number as its RC.

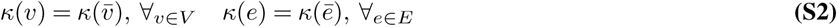

We call the combination (*G, κ*) for which *κ* satisfies Eqs. S1-S2 a *junction-balanced genome graph* (JBGG).

### Inferring JBGGs from stromally admixed tumor samples

We infer JBGGs from WGS read depth and junction data through the solution of a mixed integer quadratic program (MIQP), which assigns an integer CN *κ* : *V*_*I*_ ∪ *E* → ℕ to the vertices and edges in a genome graph *G*. The input to the optimization is a genome graph *G*, purity *α*, and ploidy *τ*. The genome graph is generated, as above, from a set of breakpoints ℬ_*seg*_ obtained from a preliminary segmentation of genome-wide read depth (i.e. via segmentation software such as CBS) and a set of junctions 𝒜 (i.e. from a junction caller such as `SvABA` or DELLY).

Every internal vertex *v* ∈ *V*_*I*_ of *G* maps to a set *X*(*v*) = {*x*}, *x* ∈ ℝ^+^ of (normalized) read depth data with mean value 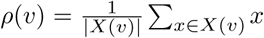. We model each data point *X*(*v*) as i.i.d. samples from a Gaussian distribution with standard deviation *σ*(*v*) and mean *µ*(*κ*(*v*)), the latter of which depends on the (latent) vertex integer clonal copy number *κ*(*v*) via an affine function *µ* : ℕ → [0, ∞). Given a tumor with purity *α* and ploidy *τ* and average genome wide read depth *ρ*_0_, we define

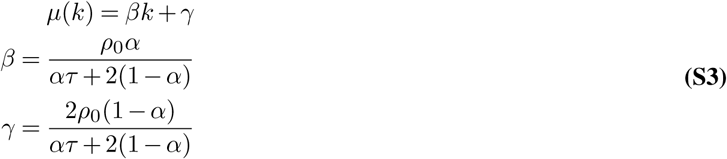

The log likelihood is

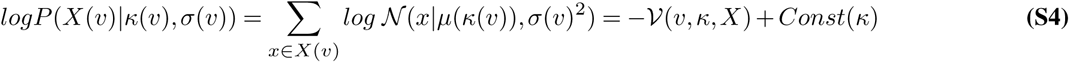

where 𝒩(*µ, σ*^2^) is the Gaussian probability density function with mean *µ* and standard deviation *σ*^2^ and 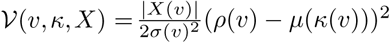 is the *read-depth residual* of vertex *v*. The standard deviation *σ*(*v*) is a *κ*-independent constant which is computed directly from the data, e.g. by taking the sample standard deviation across all vertices 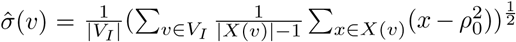. In practice, (see below) we apply more complex estimates for *σ*(*v*) to account for heteroscedasticity in the read depth data. We note that the first summand in the rightmost expression in Eq S4 is also *κ*-independent.

Given this model, the joint log-likelihood of the read depth data 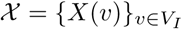 across the graph given copy number assignment *κ*:

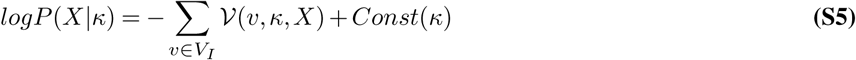

We also refer to 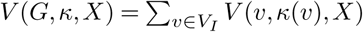 as the *read-depth residual* of the JBGG (*G, κ*) (relative to data 𝒳).

The satisfaction of junction balance and skew-symmetry constraints in Eq. S1-S2 may place nonzero copy number at one or more loose end edges. Each loose end in our graph represents a slack variable that allows the junction balance constraint to be relaxed at specific internal vertices, allowing the data to be fit when some junctions are missing from the input (e.g. due to low mappability, sequencing depth, or purity). To penalize solutions that require the use of many loose ends, we add an exponential prior with decay parameter *λ* on the loose end CN in (*G, κ*), which makes models with many missing junctions unlikely. This prior has log likelihood

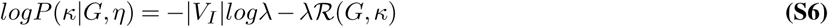

where

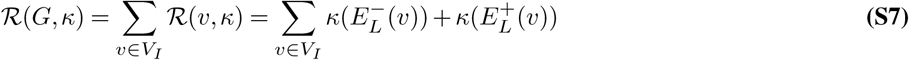

is a *complexity penalty*. Adding the log likelihood in Eq. S5 to the prior in Eq. S6 yields a penalized log likelihood for the data with regularization parameter *λ*. Under this model, the maximum a posteriori probability (MAP) estimate of *κ* will minimize the function

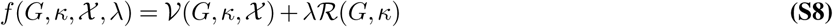

which combines the quadratic read depth residual 𝒱 and *ℓ*_1_-norm complexity penalty ℛ into a single quadratic objective. In practice, we also consider models that penalize the number of loose ends with nonzero copy number, i.e. applying an *ℓ*_0_-norm penalty 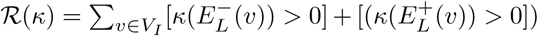. We use *f* to define a MIQP, which we solve to infer a MAP estimate for *κ* given data 𝒳 and genome graph *G*:

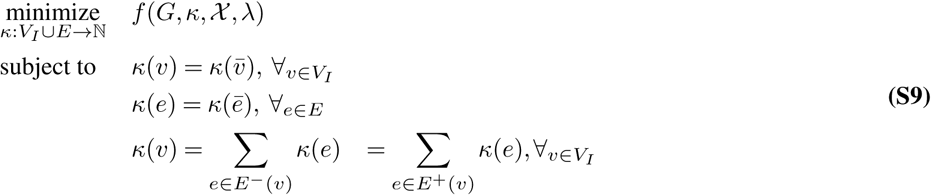

The resulting MAP estimate 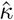 defines the JBGG 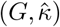 which is outputted and returned to the user.

### JaBbA pipeline

Junction Balance Analysis (`JaBbA`, https://github.com/mskilab/JaBbA) is an R package freely available under the MIT license. The only two required inputs to `JaBbA` are high density read depth data 𝒳 and a set of junctions 𝒜 (see mathematical formulation above). The key output is a `gGraph` object representing the junction-balanced genome graph, which is further queried and analyzed using the `gGnome` package (https://github.com/mskilab/gGnome). The workflow of `JaBbA` is composed of three phases: preprocessing, model fitting, and postprocessing.

### Preprocessing

During the preprocessing phase, the input coverage and junction data is transformed with **Algorithm S1** into a genome graph without copy number that 1) divides the reference genome into internal vertices *v*_*I*_ ∈ *V*_*I*_ each of which will be assumed to have a coherent CN, and 2) establishes edges *e* ∈ *E* = *E*_*R*_ ∪ *E*_*A*_ ∪ *E*_*L*_ that each represents an adjacency consistent with the reference genome (REF edge, *E*_*R*_), created by an rearrangement junction (ALT edge, *E*_*A*_), or a unmatched breakend (loose end, *E*_*L*_). The ends of the vertices are the union of junction breakpoints *getBreakpoint*(𝒜) and a primary segmentation of the genome ℬ_*seg*_ using the Circular Binary Segmentation algorithm (Olshen et al., 2004) (CBS), or from user input. At this stage, the purity *α* and ploidy *τ* are also inferred with any of the three built-in methods namely a custom least squares grid search `ppgrid` (available within `JaBbA`), package `Ppurple` (https://github.com/mskilab/Ppurple) a probabilistic purity and ploidy estimator, and package `Sequenza` (Favero et al., 2015). Purity and ploidy estimates can also be provided by user input. See below for pipeline used for the full study.

### JaBbA model fitting

With the coverage data mapped on each vertex *X*(*v*), robust mean 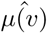 and variance 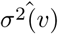 of each internal vertex *v* ∈ *V*_*I*_ in the genome graph are estimated and they determine the mapping from an integer CN state *κ*(*v*) to the read depth residual sum of squares 𝒱(*v, κ, X*) laid out in **Eq. S4**. Combining with the user defined loose end penalty *λ* (default 100, tuned for 200 bp binned tumor-normal pairs), the MIQP problem (**Eq. S9**) is solved on the genome graph. The solver is `CPLEX` (v12.6.2, IBM).

### Postprocessing

After the optimization with `CPLEX`, we further simplify the solution by merging neighboring vertices of the same total CN that are connected only through REF edges. When a normal control is present, we also annotate the allelic CN of each vertex, *κ*_*a*_(*v*), *κ*_*b*_(*v*) where *κ*_*a*_(*v*) + *κ*_*b*_(*v*) = *κ*(*v*). At each of the germline heterozygosity sites, the allelic read counts are tallied, total integer copy numbers from `JaBbA` mapped, and SNP specific ploidy is calculated as the mean of total CN across heterozygosity sites. The series of *centers* in allelic read count space corresponding to each integer somtic copy number state are then corrected the same way as described in **Eq. S3**. The integer CN state for the low count allele is then determined by maximum-likelihood under a Poisson distribution assumption of allelic read counts. The final output is then saved into a gGraph object that can be manipulated, visualized, and analyzed with the `gGnome` R package (https://github.com/mskilab/gGnome) and `gGnome.js` browser (https://github.com/mskilab/gGnome.js).

### Iterative junction rescue

By default, we recommend using the unfiltered `SvABA` candidate junction calls as the input to `JaBbA`, which will invoke an iterative fitting procedure that starts with the high-confidence calls then gradually consider extra low-confidence candidate junctions that would otherwise be filtered out by `SvABA` due to insufficient read support. Only the low-confidence junctions near (<1 kbp) a loose end from the last iteration are considered in the next, in the hope that their integration can further reduce the loss function. Such procedure is particularly beneficial to the sensitivity of the graph reconstruction when the purity, sequencing coverage fluctuate, and performs slightly better than `JaBbA.un` which uses the union of three junction callers at lower sample purities (**Fig. S1D**).

### Quality control

From the 2,833 WGS sequenced samples, we excluded 35 samples from the analysis due to these reasons: 5 did not optimize within `JaBbA` successfully, 1 failed complex event annotation due to an extremely large number of incorporated junctions, and 29 low quality samples completely failed to converge. Quality control was determined by an initial coverage-derived segmentation and junction set which yields a genome graph whose size was used as the QC metric. Initially graphs exceeding 130,000 vertices (**Fig. S5B**) were marked as low QC genomes for graph inference, since these required additional computational resources to converge (> 500 GB RAM, > 24 hours of compute time). This yielded 69 low QC genomes which `JaBbA` may not be able to resolve due to excessive computational requirements. Among these 69 hypersegmented graphs, we were able to converge to a `JaBbA` solution for 40 using additional compute resources (up to 1 week of run time). After this 2nd pass-through, we excluded 29 graphs from the remaining analyses on which `JaBbA` still failed to converge using high memory, long running jobs.

### Structural variant event classification

To identify simple and complex structural variant events in the `gGraph` output of `JaBbA`, we implemented a set of classifiers. These are implemented in the `gGnome` package (freely available from: https://github.com/mskilab/gGnome). The footprints of the discovered events on the pan-cancer dataset of 2,798 genome graphs are available in **Table S5**. Procedures followed to discover each event type were as follows.

### Rigma and simple deletions

We identified simple deletions and rigma as follows: We found groups of DEL-like junctions that were composed of at least 2 junctions with JCN < ploidy, greater than 10 kbp and less than 10 Mbp interval size. We used `fishHook` as a Poisson model on a per sample basis to statistically nominate regions of the genome enriched with DEL-like junctions using sliding windows of 1 Mbp (500 kbp stride) while correcting for the occurrence of other junction types. From the model, we chose regions that have a significant enrichment of *FDR* < 0.5. Candidate deletion clusters that pass the above filters and fall within statistically enriched regions were then marked as rigma events. Remaining isolated low-JCN deletions were called as simple deletions.

### Pyrgo and simple duplications

The procedure to identify pyrgo and simple duplications was the same as for rigma above. However, instead of DEL-like junctions, we substituted DUP-like junctions, and used the above filters. For candidate duplication clusters that pass these filters, fishHook was again used to statistically nominate regions in a genome enriched for duplications while correcting for the occurrence of non-duplication junctions. Candidate duplication clusters that fall within the statistically enriched windows were then marked as a pyrgo. Remaining isolated low-JCN duplications were called as simple duplications.

### DM, BFBC, and Tyfonas

Across 2,798 `JaBbA` graphs, to nominate amplicons (amplified clusters) weakly connected components were identified within `JaBbA graphs` amongst segments (i.e. vertices) having a minimum segmental copy number of > 2×ploidy. 12,327 subgraphs were nominated from this initial clustering which were further subsetted to only those that contained a maximum JCN > 7, leaving 1,675 high level amplicons with a high copy JCN. Four curated features were used to annotate each of these amplicons, 1) maximum segmental/interval copy number (MICN), 2) the sum of fold back inversion JCN normalized by MICN (FBIJCN), 3) maximum JCN normalized by MICN (MJCN), and 4) the number of high copy junctions (thresholded on JCN≥ploidy), NHIGH. To separate these amplicons, hierarchical clustering was performed upon these features using Euclidean distance and complete linkage, cutting the tree to arrive at *k* = 3, with each group corresponding to patterns of double minute (DM), breakage fusion bridge cycle (BFBC), and tyfonas events.

A decision tree was trained using recursive partitioning (function `rpart` in the eponymous R package) from the hierarchical clustering result based on the above features. The decision tree arrived at an FBIJCN ≥ 0.61 to distinguish tyfonas/BFBC amplicons from the DM amplicons. Out of the amplicons with FBIJCN ≥ 0.61, amplicons with NHIGH ≥ 27 were called tyfonas and the rest were called BFBC. This algorithm was then used to nominate the tyfonas, DM, and BFBC from high level amplicons identified within this study. These decision tree-based calls were then used for the subsequent analyses on these amplification events (see below).

### Chromothripsis

For chromothripsis we searched for the subgraphs that best fit the features of chromothripsis patterning described in (Korbel and Campbell, 2013). Candidate clusters within the graph (i.e. weakly connected components) were nominated that contain a high density of overlapping junctions, and whose segmental copy numbers occupied a narrow range of states. Specifically, each vertex within every footprint of a candidate cluster was subjected to three criteria: 1) occupancy of at most 3 different copy states (for a diploid genome, adjusted up proportionally to the ploidy of the genome), 2) composition of at least 8 segments, and 3) having an interval copy number at the width-weighted 99^th^ percentile that did not exceed 4 (for a diploid genome, or adjusted to the ploidy of the genome). Clusters that survived this three-step filter were required to contain junctions with a near-uniform mixture of basic orientations (*χ*^2^ test, p-value > 0.001) as a result of the expected random generation of junctions during chromothripsis. Out of the final remaining clusters, several further criteria were required to yield the final chromothripsis calls as follows: each cluster must have 1) at least 7 internal junctions, 2) no fewer than two sub-100kbp footprints and no more than four ≥ 100 kbp segments within the cluster, and 3) on average at least 3 junctions that overlap one another.

### Chromoplexy

Chromoplexy as first described in (Baca et al., 2013) can be identified by a series of reciprocal (or nearly-reciprocal, with small deletion bridges between adjacent breakpoints) long-range junctions. Accordingly, chromoplexy was identified from a pool of low-JCN (≤ 3) edge clusters in which junction breakpoints are no further than 10 kbp away from the next junction breakpoint. Edge clusters that contained at least 3 junctions that span more than 10 Mbp on the reference, and whose footprints occupied at least 3 discontiguous genomic territories separated by >10 Mbp on the reference were called as chomoplexy events.

### Templated insertion chains (TICs)

We nominated templated insertion chain (TIC) events that result from short insertions that involve more than 1 junction, usually resulting in a gain of copy of all loci within the event, or linking disparate loci through shorter segmental “hops” within the genome. This likely is the result of replicative error mechanism, involving template switching, which parsimoniously explains the resultant rearrangement topology (Li et al., 2017). To capture this, the following procedure was used. Breakpoints identified across the whole genome were ordered onto reference coordinates, and pairs of breakpoints that fall within an interval of ≤ 500 kbp and ≥ 50 kbp of each other were kept as candidates. These candidate breakpoint pairs must have contained 2 breakpoints of +,− (i.e. right-facing, left-facing) orientation when reading from left to right on reference coordinates with each breakpoint originating from 2 separate junctions. A new graph with vertices comprising each candidate junction pair as vertices and edges comprising connections between junction pairs (linked by short candidate intervals between breakpoints) was constructed. All paths (i.e. walks) through the graph that traverse through at least 2 vertices/junctions were obtained. ALT junctions within each walk traversed on this graph were then labeled as unique TIC events.

### Simple translocations, inverted duplications, and inversions

We called inversions (inv), inverted duplications (invdup), and translocations (tra). Inversions and inverted duplications were both defined as pairs of overlapping, oppositely oriented, INV-like junctions of the same JCN as well as equal left and right vertex CN, with no third junction or a loose end interfering. We defined translocations as single or reciprocal pairs of TRA-like junctions connecting two different reference chromosomes that are not connected by any other junctions.

### gGnome.js visualization portal

To visualize the genome graphs we developed `gGnome.js` (source code at https://github.com/mskilab/gGnome.js), a Javascript based genome browser that draws genome rearrangement graphs and allows for interactive browsing in any number of discontinuous genomic intervals accompanied by various types of genomic annotations. The flexibility of arbitrary window browsing provided crucial support for complex rearrangement events. The annotations of events were indexed within the browser and can be searched and presented. Thanks to the interface with `gGnome` we could visualize any genome graph, like the ones whose CN is inferred by other graph callers, or even plain genome graphs without CN.

### Cancer genome analysis

Harmonized pipelines for junction detection, high-density read depth estimation, purity / ploidy inference, somatic SNV / indel calling, and loss of function mutation calling were applied to the full dataset of 2,833 WGS cases as described below. Several standard analyses for mutational interpretation, signature inference, and APOBEC analyses were performed.

### Junction detection

Somatic structural variant (SV) calls were obtained using `SvABA` (Wala et al., 2018), without filtering, based on the optimal settings for input into `JaBbA` (see above). To eliminate germline artifacts, we additionally obtained germline SVs available through `SvABA` and constructed a panel of normals (PON) consisting of all identified germline SV across the 1,017 TCGA tumor/normal pairs. All germline SVs were de-duplicated by overlapping every pair of SV breakpoints within a 1,000 base pair window. All sets of overlapping SVs were collapsed into one unique SV. Somatic SV calls were then filtered through this PON. For the CCLE cell lines for which a paired normal does not exist, we used standard WGS from HCC1143BL as the normal reference sample to call SVs. Germline SVs were then filtered using the above described PON.

### Read depth

We obtained a coverage profile for every BAM alignment in our cohort, extracting read counts for every 200 base pair tile across hg19 reference coordinates via `samtools` (v1.9) and adjusted the tile-level read counts by locally estimated scatterplot smoothing (function `loess` within R) fitting them against the mappability score and GC proportions for each tile as covariates. We then obtained an initial copy segmentation profile by inputting paired tumor coverages to corresponding normal coverages into a modified implementation of the Circular Binary Segmentation algorithm (Olshen et al., 2004). As a normal coverage profile for the CCLE cases, we used a composite of the 1,017 TCGA normal coverage profiles comprised of the average of the 200 bp bins across all autosomal chromosome coordinates. For bins on X and Y chromosomes, we doubled the X and Y coverage from males and then averaged the X and Y coverages from males and females combined to create a uniform coverage profile serving as a universal normal for the unpaired CCLE samples.

### Purity and ploidy estimation

For all cases with a tumor and normal pair, we obtained germline heterozygous SNP allele counts by intersecting SNV sites present in both tumor and normal with SNPs from HapMap 3.3. We obtained purity and ploidy estimates for all samples through `Sequenza` (Favero et al., 2015), `TITAN` (Ha et al., 2014) or a custom least squares grid search `ppgrid`, (available through `R` package, `JaBbA`). We then used the consensus purity and ploidy from the panel of the three calls across all tumor normal pairs. For CCLE cell lines we only used `ppgrid` to obtain purity and ploidy estimates since germline SNP sites were unavailable.

### Somatic SNV and indels

To obtain somatic SNV/InDel calls `Strelka2` (Kim et al., 2018) was run under paired tumor and normal mode with default parameters using hg19-based references. Somatic SNV and indels were obtained only for those cases where tumor and normal BAMs were available (2494 out of 2833 cases). In addition to the recommended filters, a universal mask was used to remove common artifacts in low-mappability regions described in (Mallick et al., 2016). After these initial filters, only sites determined to pass `Strelka2`’s quality filter (i.e. sites where the “FILTER” field was marked as “PASS”) were considered, yielding a high quality set of somatic SNV calls. Variant annotations were obtained for SNV/InDel using SnpEff with the GRCh37.75 database.

### Germline SNV and indels

For germline variants, SNV/InDel calls `Strelka2` (Kim et al., 2018) was run as above, except in normal-only mode. Germline SNV and indels were obtained only for non-cell line samples within the study. The universal mask was also used for germline SNV/InDel calls. Additional filters restricted the germline variants used to those which met the following criteria: i) sites that do not overlap with common variants i.e. those variants that matched coordinates and ALT alleles with sites from the normal ExAC population that have a minor allele frequency of >1% (ftp://ftp.broadinstitute.org/pub/ExAC_release/release0.3.1/subsets/), ii) high quality sites as determined by `Strelka2`’s quality thresholds (i.e. sites in which the “FILTER” field was marked as “PASS”), and iii) sites that overlapped and matched ALT alleles with known pathogenic variants from ClinVar annotations ftp.ncbi.nlm.nih.gov:/pub/clinvar/vcf_GRCh37. Variant annotations were obtained for this final, high quality set of germline SNVs/InDels using SnpEff as above.

### Mutation interpretation

For alteration status of genes across the cohort we obtained loss-of-heterozygosity calls using allele-specific copy number available for cases with tumor/normal pairs using germline heterozygosity coverage genome-wide for these samples. Homozygous deletions, heterozygous deletions, and amplifications were detected from absolute copy number calls available from the `JaBbA` graphs. Filtered SNV/InDel annotations falling within protein coding regions, were considered only if they constituted somatic missense or truncating events, germline pathogenic variants, or germline truncating variants. For CCLE, our callset only included the publicly available mutational drivers. For testing mutational status, only those DNA damage genes which intersected between Cancer Gene Census (CGC) (Sondka et al., 2018) genes and genes in DNA damage repair pathways as annotated in (Pearl et al., 2015) were used. Final lists of genes that were nominated significant and mutational status within those genes per sample are found in **Table S2**.

### SNV signatures

To obtain contributions of the 30 COSMIC (https://cancer.sanger.ac.uk/cosmic) mutational signatures (release 2) (described in ref. (Alexandrov et al., 2013; Nik-Zainal et al., 2012; Tate et al., 2018)) within all tumor/normal pairs, we employed `deconstructSigs` using the high-quality somatic SNV calls as input (Rosenthal et al., 2016). Only SNVs falling within well-mappable sites in the human genome were considered, i.e. those that occur outside of the universal mask regions described above. Genome-wide signature weights were corrected using the ratio of trinucleotide frequencies in the hg19 genome over the ratio of the trinucleotide frequencies in the eligible regions. The re-weighted 30 COSMIC signatures were refitted to the cohort to obtain signature contributions. The most likely SNV signature was determined for each somatic SNV call within a sample by using the vector of fitted signature contributions as the prior multiplied by the vector of signature weights possible for a given trinucleotide context. To obtain the likelihood of a variant arising from each of the 30 signatures given the signature contribution, each trinucleotide context weight was then normalized by the sum of the product of the signature contributions and trinucleotide signature weights. All SNVs whose maximal likelihood corresponded to Signature 2 or Signature 13 were considered as APOBEC driven SNV. All others were non-APOBEC driven.

### APOBEC mutations

We additionally defined APOBEC as GC-strand coordinated clusters originally described in (Roberts et al., 2013) by fitting the distance between SNVs on the reference (distance defined per sample) to an exponential model (glm function from the stats package within R with argument “family = Gamma”, and with a shape parameter of 1 to score distance between breakpoints). The model was fit using all inter-breakpoint distances to get a dataset-wide average rate. P-values were derived by comparing the observed inter SNV distances to the expected using the fitted parameters based on the exponential generalized linear model. All breakpoints with an *FDR* < 0.10 were considered clustered. Clusters were then identified by labeling consecutive runs of clusters on the reference. Consecutive runs with only G or with only C mutated were labeled as GC-strand coordinated.

### Genome simulations

For the data to benchmark `JaBbA` genome graph reconstructions, we synthesized two series of WGS data. First we generated a set of *de novo* forward simulated, rearranged cancer genomic sequences from an initial set of input junctions (`SimBLE`, https://github.com/mskilab/sim.ble). SimBLE iterates through simulated cell cycles to gradually incorporate the input junctions into the derived genome from previous steps until exhausting the input junction set, while keeping track of the actual rearranged haplotypes. In the end, it generates a coherent FASTA file of the rearranged genome guided by the haplotypes encoded in lists of reference genomic ranges. We then simulated sequencing reads from this FASTA file with ART read simulation software (Huang et al., 2012) to an average depth of 40X and aligned them to the reference genome hg19 to obtain the simulated BAMs. Trivially, the reference genome itself is also subjected to the same *in silico* sequencing to provide as normal controls. We did 40 distinct simulations with different subsets of varying size extracted from the junctions identified in HCC1143 breast cancer genome. In addition to these simulated BAMs, we also obtained WGS for HCC1954 breast cancer cell line and HCC1954BL the corresponding normal fibroblast cell line. Finally, we take these BAM files together, downsampled and mixed reads from matching tumor and normal to a series of ten tumor read proportions, from 0.1, to 1.0, mimicking the stromal cell admixture of real tumor samples. We created four technical replicates at each of the ten purity levels, and ended up with 40 pairs of tumor and normal BAM files for the 40 distinct simulated genomes and HCC1954, respectively.

### Benchmarking JaBbA

In the simplest terms, reconstructing genome graphs consists of two tasks: estimating junction copy numbers and DNA segment copy numbers. A junction is said to be incorporated in the genome graph if it is assigned non-zero copy number. Thus, a genome graph reconstruction method’s performance can be evaluated from these two main aspects, 1) incoporating correct junctions and estimating the exact JCNs, and 2) faithfully segmenting the genome and estimating CNAs. We compared `JaBbA`’s performance in both aspects against three other genome graph reconstruction methods (`PREGO`, `Weaver`, and `ReMixT`), and specifically in segmental CN estimation with five extra somatic CNA callers that do not infer genome graphs (`BIC-seq`, `FACETS`, `TITAN`, `FREEC`, `CONSERTING`).

To make a fair comparison, all four genome graph reconstruction methods received harmonized junction inputs consisting of the union of `SvABA`, `Delly`, and `Novobreak` (“union junction set”) as input junctions. Additionally, `JaBbA` and `Weaver` also ran with their respective default recommendations, `JaBbA`’s being `SvABA` unfiltered candidates and `Weaver`’s its internal junction caller. To differentiate these two extra settings, in **Fig.S1C-H**, the version of `JaBbA` and `Weaver` with union junction set input were labeled as `JaBbA.un` and `Weaver.un`.

For FACETS and TITAN inputs we computed heterozygous SNP counts at 4,165,754 common SNP sites (GATK bundle hapmap 3.3.b37.vcf.gz, https://software.broadinstitute.org/gatk/download/bundle) using the `R / Bioconductor` package `Rsamtools`. Notably, for FACETS we used all the sites as the input instead of the usual heterozygous-only set, after consulting with the authors.

### Junction incorporation and JCN estimation

To evaluate junction incorporation and JCN estimation, the incorporated junctions were approximately matched to gold standard junctions, e.g. both breakpoints were within 1 kbp from the corresponding gold standard junctions with the same orientations. Naturally, true positives were the number of matches, false positives were incorporated junctions without a match, and false negatives were gold standard junctions without a match. F1 score of the incorporated junctions, the harmonic mean of precision and recall, was then computed for each benchmarking sample (**Fig.S1D**). Gold standard junctions for HCC1954 were defined as the consensus set inferred from the original data (junctions identified by at least two different callers).

While in real samples the true JCNs are often unknown, the simulated genomes provided us with an opportunity to directly compare fitted JCN to the ground truth. So, after the padded matching (described above), inferred JCNs of the graph reconstruction were compared to the matching truth junctions, if any, in the simulated dataset. The proportion of correctly fitted JCNs out of all incorporated junctions times recall of incorporation represented the accuracy or completeness of JCN estimation (**Fig.S1C**).

### Segmental CNA estimation

When evaluating CNA inference performance, two metrics were considered. One was the correct placement of CN change-points. Too many (hypersegmentation), too few (undersegmentation), too distant from the gold standard (large error margin) were all adverse indications. Analogous to matching junction breakpoints, we considered an inferred CN change-point to match a gold standard if their distance was within 1 kbp and they have the same direction of CN change, e.g. increasing or decreasing CN from the side with smaller coordinates to the larger. To prevent extreme cases of hypersegmentation being over-optimistic, each one of the gold standard CN change-point was allowed have at most one match (the closest one). Based on the matching, F1 scores computed and shown in **Fig.S1E**. For HCC1954, the gold standard CN change-points were defined as the consensus junction breakpoints.

The other metric was the concordance between the inferred CN profile with the gold standard. For each segment in the gold standard, the overlapping portions of all the inferred segments were identified. The inferred CN for that gold standard segment was defined as the overlap width-weighted average of inferred CN. Subsequently, the Spearman correlation coefficient was computed across all gold standard segments **Fig.S1F**. The gold standard for HCC1954 CN profile was derived from array-CGH downloaded from the CCLE data portal.

### Validation of junction copy numbers with long-range sequencing technologies

To show that the `JaBbA` inferred junction copy numbers are accurate in real samples, we used 10X linked-read sequencing (10X) of the breast cancer cell line HCC1954 as orthogonal experimental validations. At a set of selected junctions of various JCNs, the number of unique barcodes mapped within 20 kbp from both *cis* sides of the breakpoints of each ALT junction was plotted as a red dot in **Fig.1C**. Same applied to a range of REF junctions (blue dots). Finally, both were compared to the fitted ratio between barcode coverage and DNA integer CN at randomly sampled unrearranged loci (grey line).

### Analysis of reference junction clusters

We defined clusters of low JCN junctions as groups of at least 3 junctions with JCN < ploidy whose intervals all cross either between the breakpoints (for intrachromosomal junctions) or within a 100 kbp window from interchromosomal breakpoints. We analyzed the density of low-JCN DEL-like or DUP-like junctions across the genome within our cohort using a gamma-Poisson model, `fishHook`, previously described in (Imielinski et al., 2017) to identify loci that are significantly enriched with the desired genomic event while accounting for other junctions which may also be enriched concomitantly.

Specifically, across 1 Mbp sliding windows (500 kbp stride) as the response, we counted the number of low-JCN deletions or duplications within each sample and ascertained bins that have higher density of events than expected under the model taking all bins across all samples and the local density of non-deletion or non-duplication junctions, respectively, into account. P-values were obtained for every tile × sample combination by comparing the observed density of each tile/sample pair against the expected value of the gamma-Poisson model. Tiles that were density outliers by an FDR threshold < 0.10 were considered density outliers for visualization. Based on this model fitting, we defined algorithms to search for pyrgo and rigma events on `JaBbA` graphs (see above for precise procedures).

### Event topography

Several sources of chromatin data were used to determine topographic correlations between event (rigma, pyrgo, simple deletion, simple duplication) density and genomic covariates. Smoothed replication timing data from lymphoblastoid cell lines (personal communication with Amnon Koren) was used as the reference to determine early, middle, and late replicating regions in the genome. For superenhancer analysis, we considered the union of coordinates from all cell lines with superenhancer peaks described in (Hnisz et al., 2013). Human fragile sites were obtained from querying NCBI (https://www.ncbi.nlm.nih.gov) gene databases. Gene annotations were obtained from GENCODE v29 lifted to hg19 coordinates.

### Rigma timing in BE

To assess the timing of rigma and deletions, analysis was restricted to the 347 samples from 80 cases for which we had multiple biopsies (i.e. the Barrett’s esophagus cohort). All genomic footprints of rigma and simple deletion events were analyzed for overlap with other events across samples for each case. Deletions or rigma which were found within a 500 bp window in at least one of the other samples for an individual case were designated as early.

### Comparison of rigma and chromothripsis

To compare chromothripsis and rigma events, we assessed two different measures. First was the genomic footprint occupied by each event type. Second was the number of junctions that could maximally be placed in *cis*. This measure was obtained by seeking the longest possible (traversing through the most junctions) allelic haplotypes through the `JaBbA` graphs (as defined above), through the loci containing these two events. Junctions that occurred on one haplotype in a given locus were considered in *cis* whereas those occurring across different haplotypes were in *trans*.

### Local allelic reconstruction with linked-read sequencing

Within a locus of interest, we used 10X linked-read sequencing to reconstruct the set of local allelic haplotypes that best represents the sequencing data while conforming to the copy number constraints of the local graph. Potential haplotypes *H* are enumerated by identifying all traversals through the local subgraph *G* from source to sink vertices. An allelic haplotype *h* ∈ *H* is a walk through subgraph *G* containing a sequence of vertices *V* (*h*) ⊆ *V* (*G*) and edges *E*(*h*) ⊆ *E*(*G*). For every linked read *p* from a linked-read dataset 𝒫 and every haplotype *h* ∈ *H*, the function *O*(*p, h*) → {0, 1} indicates whether the linked read *p* supports the haplotype *h*. A linked read (a set of aligned reads from the 10X linked-read sequencing library that share a barcode) supports a given allelic haplotype *h* (thus, *O*(*p, h*) = 1) if it meets all of the following conditions: (1) all of the individual reads from the linked read with alignments within the window of the locus must align to vertices included in the allelic haplotype *v* ∈ *V* (*h*); (2) the vertices *v* ∈ *V* (*h*) with linked-read coverage must be consecutive within the walk *h*; and (3) if *h* contains any ALT junction *e* = (*v*_1_, *v*_2_) ∈ *E*_*A*_, the linked read must have alignments to both *v*_1_ and *v*_2_ (i.e. junction-supporting reads) for at least one junction in the allelic haplotype. A mixed integer program (MIP) assigns a copy number to each allelic haplotype subject to the constraints of the local vertex and edge copy numbers. The objective function minimizes the number of unique haplotypes *h* ∈ *H* assigned a non-zero copy number while maximizing the number of linked reads with support for at least one *h* ∈ *H* assigned a non-zero copy number. The MIP is defined as follows:

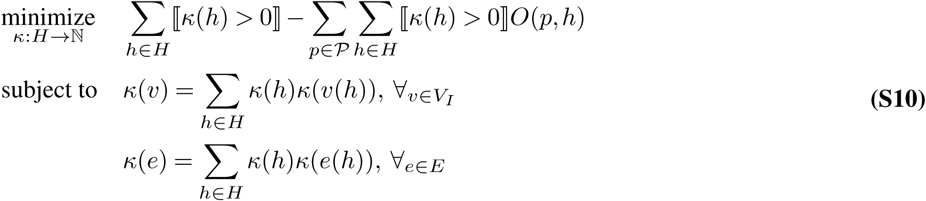

where the definition of *κ*, defined previously as a mapping *κ* : {*V*_*I*_ ∪ *E}* → ℕ of non-negative integer copy number to vertices and edges of *G*, has been expanded to *κ* : {*V*_*I*_ ∪ *E* ∪ *H}* → N, where *κ*(*h*), *h* ∈ *H* represents the non-negative integer copy number of a given allelic haplotype *h*, and *κ*(*v*(*h*)) and *κ*(*e*(*h*)) represent the copy number within a single copy of *h* of vertex *v* and edge *e* respectively.

### Max *cis* analysis

To characterize the *cis*-allelic structure associated with select complex event patterns (i.e. rigma and chromothripsis) without explicitly inferring allelic phases, we developed an optimization approach to search through all possible allelic states to identify the configuration that placed the most ALT junctions in *cis*. We apply this “max *cis* analysis” to each local genome subgraph defined around a given event. The weight *w* of a single flow *f* ⊆ *E*(*G*) through the graph from a source vertex to a sink vertex is defined as *w*(*f*) = Σ_*e*∈*f*_ ⟦*e* ∈ *E*_*A*_⟧, such that *w*(*f*) is equal to the number of ALT junctions in *f*. The flow that incorporates the maximum possible number of ALT junctions is found by solving a MIP with the following objective function:

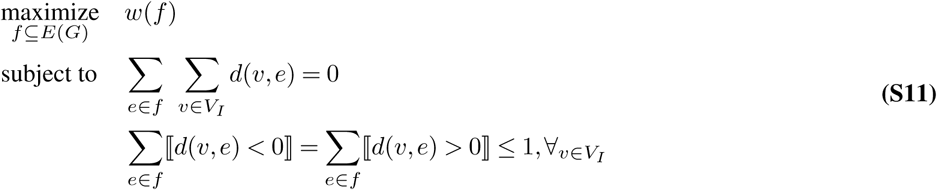

where *d* is defined for a vertex *v* and edge *e* as

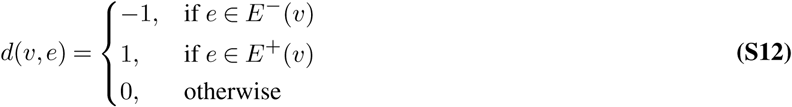

The linear allelic haplotype *h*_*f*_ corresponding to flow *f* is the walk through the local subgraph G that satisfies *E*(*h*_*f*_) = *f* ⊆ *E*(*G*). Because some events include tandem duplication or fold back junctions that create cycles on the graph, it is also necessary to specifically consider a potential cyclic allele *ĥ*. We do this by searching the subgraph for clusters of vertices which are strongly connected components. We define *ĥ* as the largest subset of strongly connected vertices *V* (*ĥ*) ⊆ *V* (*G*). If *h*_*f*_ and *ĥ* share any vertices, we define *f*_*T*_ as the combination of *f* ∪ *E*(*ĥ*). Otherwise, *f*_*T*_ is either *f* or the edges associated with cyclic allele *ĥ* based on which contains a greater number of ALT junctions.

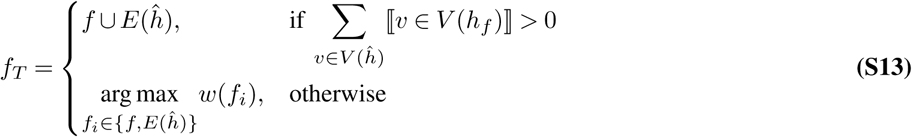

The fraction of ALT junctions in *cis* is the number of ALT junctions incorporated into *f*_*T*_ over the total number of ALT junctions in the local subgraph, or:

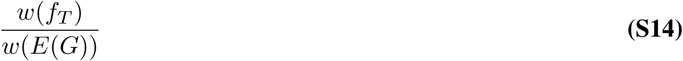

### Driver amplification analysis

Analysis of amplification driver genes falling within complex amplification events as defined above (DM, BFBC, tyfonas) employed GISTIC amplification peaks publicly available from (Zack et al., 2013), which intersect with Cancer Gene Census genes. Any overlap between the genomic footprint of a DM, BFBC, or tyfonas event with an amplification driver was counted to calculate the frequency of each events’ incidence upon one of these genes.

### Kataegis

To analyze the phenomenon of kataegis incident near rearrangements from complex amplicons, all SV breakpoints were collapsed onto a new coordinate system with a single axis in which coordinate 0 corresponds to the location where the breakpoints occur. Positive coordinates point towards segments that are involved in the rearrangement (*cis*) and negative coordinates involve coordinates that are uninvolved in the rearrangement (*trans*). Only pairings of one SV breakpoint to its nearest SNV within a case are matched, and due to possible artifacts, those SNVs within 10 base pairs of a rearrangement were removed for each sample. The relative density of SNVs at every base pair on the new coordinate system was calculated by dividing the total number of SNVs at every site by the average the number of SNVs from 0 kbp to −5 kbp (i.e. SNVs on the uninvolved segments from 0 kbp to 5 kbp away from rearrangement breakpoints). APOBEC driven kataegis is defined using SNV using the footprints of APOBEC activity as defined above (SNV attributed to COSMIC SNV signatures 2 and 13, GC-strand coordinated clusters, or C mutations in occurring within the TpC context).

### Clustering pan-cancer samples by SV event burden

The total count of junctions attributed to each of the 13 described event types (3 novel events, 5 known complex events, and 5 simple events-see above for calling procedures) were used as features to hierarchically cluster the data cohort. The counts for those cases with multiple samples were averaged across each feature to obtain a mean value for those samples. Each event was normalized as a negative-binomial-(or gamma-Poisson-) distributed z-score (implemented in the edgeR R package). The *µ* and size parameters of each event burden distribution is estimated using maximum-likelihood (implemented in the fitdistplus R package). Hierachical clustering upon the sample-by-normalized count matrix was performed using the Euclidean distance metric and the “Ward.D2” agglomeration method.

### Survival analysis

We obtained survival data for 1,767 cases in our cohort from cBioPortal (https://www.cbioportal.org/) for TCGA cases and from the for TCGA cases and from the ICGC Data Portal (https://dcc.icgc.org) for ICGC cases. Kaplan-Meier analysis was performed, with the sample clusters stratified above compared to the Quiet cluster using the `survival` and `survminer` R packages. Additionally, survival was compared between each of these cluster groups to the survival of those within the Quiet cluster by performing Cox regression to correct for covariates that could impact survival due to the heterogeneity of features within each cluster. The following variables were included in the model: age, sex, tumor type, status of TP53 loss of function, metastasis, tumor mutational burden, and rearrangement burden. Age, tumor mutational burden, and rearrangement burden were encoded as binary covariates, by stratifying the data points corresponding to the variables into 2 quantiled groups each: below the upper 3rd quartile (75th percentile), and above the upper 3rd quartile.

### Statistical tests

Generalized linear modeling within `R` (version 3.5.1) was used for significance testing where noted. For binary outcomes, standard logistic regression or, wherever noted, a bayesian implementation of logistic regression (available through the `arm` package within `R`, function `bayesglm`, with argument `family = “binomial”` was used. Ordinal logistic regression in **Fig. 2** (available from `R` package `MASS`, function `polr`) was used as well for testing replication timing as an ordered response variable. For count data, gamma-Poisson regression (`MASS` package, function `glm.nb`) was used. For gamma-Poisson and logistic generalized linear models, Wald tests were used to determine significance of parameters. Gamma-Poisson statistical modeling procedure (`fishHook`) to nominate pyrgo and rigma is described above (also, see (Imielinski et al., 2017)).

To statistically test for the enrichment of kataegis specifically, a gamma-Poisson regression model was employed with the response being the number of mutation counts within the *cis* territory or *trans* territory of every breakpoint. The categorical variable of whether the SNV counts occur in *cis* or *trans* territory was included to estimate the enrichment of *cish*-occurring over *trans*-occurring SNVs. Finally within the model an interaction term was included between the variable of whether the territory corresponds to a breakpoint of a DM, BFBC, tyfonas, or other event type, and the *cis/trans* categorical variable. This interaction term can be interpreted as the additional effect that an event produces with respect to the *cis/trans* enrichment (i.e. the change in kataegis that an event produces).

Where stated, Fisher’s exact test was done using `fisher.test` within R. FDR or Bonferroni correction was performed where noted to nominate statistically significant associations (`p.adjust` within R). Bonferroni cutoffs of < 0.05 and FDR cutoffs of < 0.10 were applied for each correction method where group comparisons were performed. Also, wherever stated, Wilcoxon rank-sum test was used for non-parametric testing between two groups (function `wilcox.test`, within `R`).

All p-values except for those reported in Q-Q plots are reported with a lower limit of 2.2 × 10^−16^.

### Software used

For all analyses, we used `R` (3.5.1) with `Bioconductor` (3.8, https://bioconductor.org/news/bioc_3_8_release/). Specific `R/Bioconductor` packages used in these anlayses include: `GenomicRanges` for manipulation of genomic intervals, `ComplexHeatmap` for heatmap data, `dplyr` and `data.table` for tabular operations. In addition, we employed custom Imielinski Lab packages `gUtils`, `gTrack`, `bamUtils`, and `gGnome` for manipulation and visualization of genomic ranges, sequence data access, and genome graph queries. `JaBbA` employs the IBM `CPLEX` MIQP optimizer, which is available under academic licensing (https://www.ibm.com/analytics/cplex-optimizer). For Cox regression analysis, we used `survival`, and `survminer` packages. All graphs with the exception of track data, Kaplan Meier curves, Cox hazard ratio plots, Q-Q plots, genome graph vertex illustrations, and heatmaps were plotted using `ggplot2`. Track data were plotted using `gTrack`. Decision trees were trained using `rpart`. Reordering of hierarchical cluster dendrograms was done using package `vegan`. Genome graph illustrations were plotted using the R API of `igraph`. For all additional operations to manipulate genomic/ranged data, please see the authors’ laboratory github (https://github.com/mskilab) page to access R packages: `gGnome`, `gUtils`, and `gTrack`. All other dependencies within these R packages are available from https://cran.r-project.org/ or Bioconductor R repositories.

## Supplementary tables

**Table S1. Sources of pan-cancer WGS datasets used in this study.** An Microsoft Excel spreadsheet in .*xlsx* format, summarizing of the number of samples from each dataset, the completeness of the pipelines, and the reference if the dataset is published before.

**Table S2. Sample specific features.** A *comma separated values* format text file of all the sample specific features of the pan-cancer cohort used in this study. The unique identifier for each sample and patient is stored in the *sample id* and *participant id* columns, respectively. The columns whose names start with an event type, represented the number of events (*count*), the total junctions attributed to that event type (*burden*), and the normalized zscore among the cohort used in the clustering (*zscore*). The membership to one of the 13 clusters (**Fig. 5A**) is listed under *cluster* column. It also records the gene mutation status, tumor types and subtypes, SNV density and signatures, as well as overall survival (OS) data.

**Table S3.** `JaBbA` **output segmentation** A BED file, where the first three columns of each row define a segment’s chromosome name, start coordinate, and end coordinate of a genomic range. The following columns store the metadata including sample id, total CN, event footprints.

**Table S4.** `JaBbA` **output junctions.** A BEDPE file, where the first six columns of each row define the chromosome name, start coordinate, and end coordinate of two breakpoints of a junction, and the seventh and eighth columns the orientations, each row in this file represents a junction. Annotated in the rest of the columns including sample id, JCN, and event annotation.

**Table S5. Simplified** `JaBbA` **event footprint** While Table S3 stores the complete information of all segments, this simplified CSV file contains only the locations of the footprints of annotated events.

## Supplementary figures

**Fig. S1.**
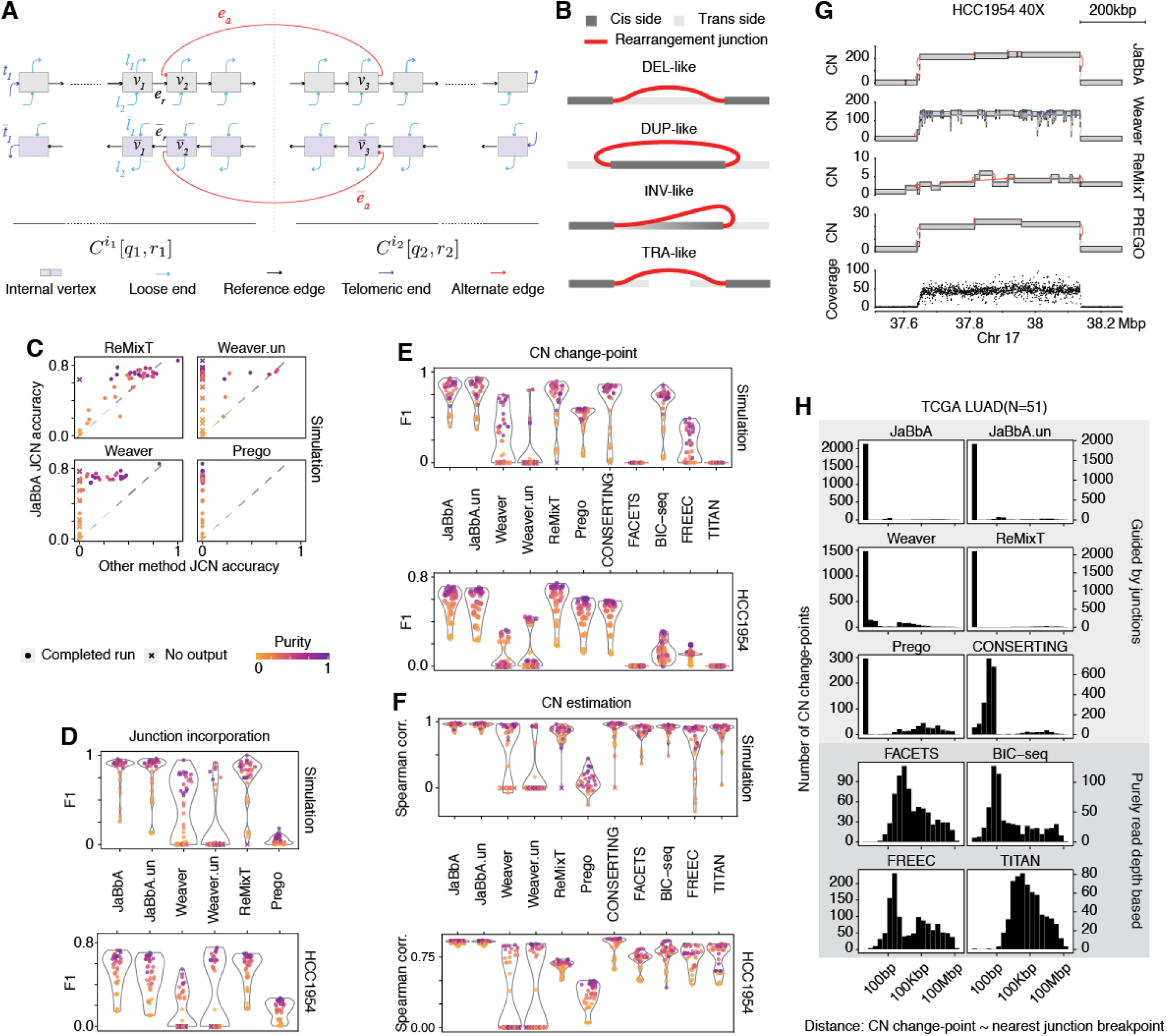
Benchmarking of JaBbA. (A) Definition of a genome graph. (B) Four basic orientations of junctions, darker grey indicates *cis* side and lighted grey *trans* side of a breakpoint. (C-F) Benchmarking in admixed simulated genomes and HCC1954. `JaBbA.un`, `Weaver.un`: `JaBbA` and Weaver with union junction as input. Unsuccessful runs with no complete output is marked as a cross. F1 score is the harmonic mean of precision and recall. (C) Accuracy of JCN estimation, (D) F1 score of incorporated junctions, (E) F1 score of inferred CN change-points, (F) concordance of inferred segmental CN with gold standard or orthogonal technology. (G) Example putative BFBC locus in HCC1954 and the reconstruction by four different methods. (H) Distances from estimated CN change-points to the nearest consensus junction breakpoints in 51 TCGA lung adenocarcinoma genomes.

**Fig. S2.**
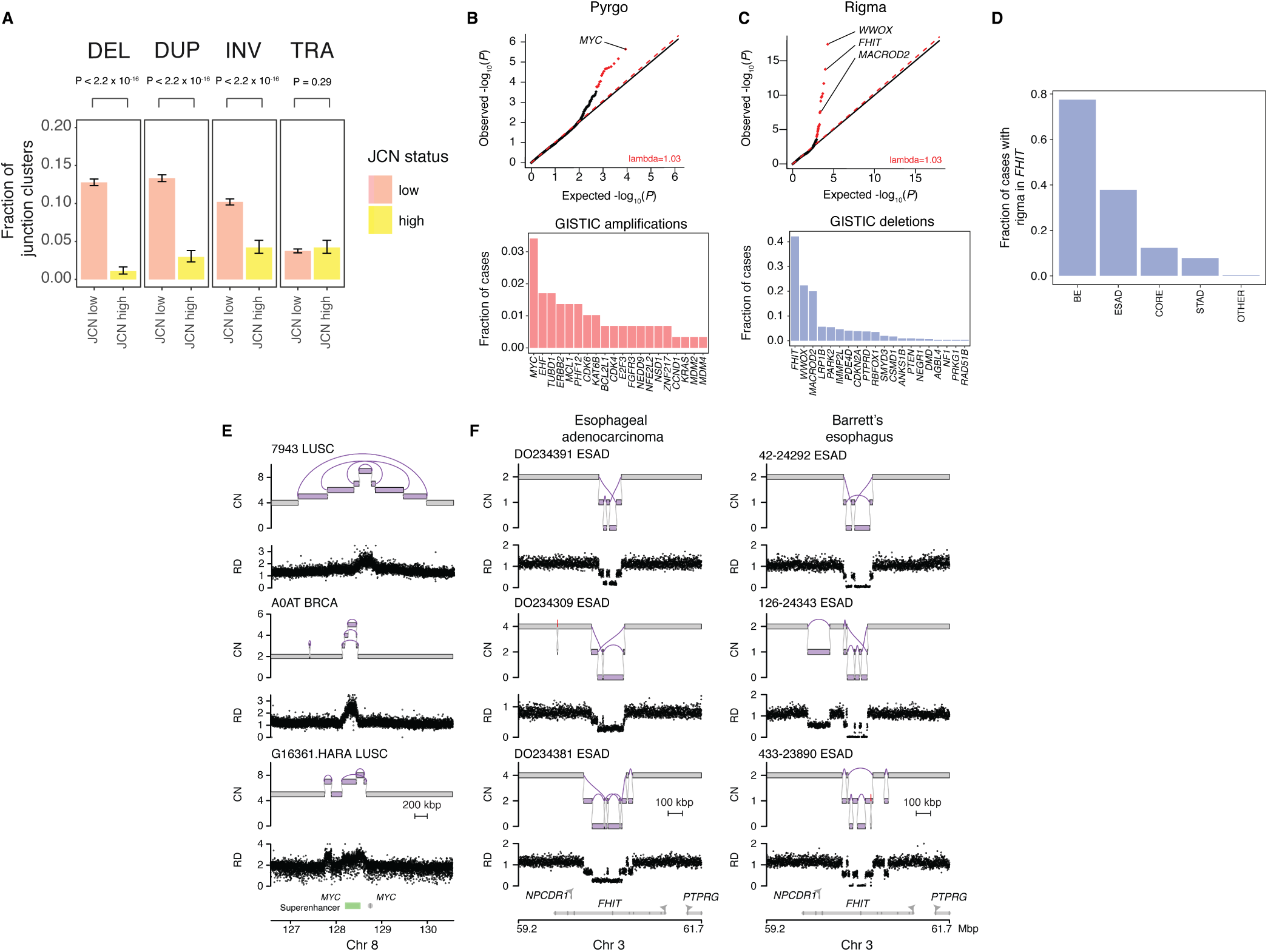
Rigma and pyrgo are specific processes involving deletion and duplication, which both selectively target cancer-related genes. (A) Clusters of junctions tend to be more homogeneous (dominated by a single junction class) at low copy number than at high copy number. Clusters of junctions are composed of at least 3 overlapping intervals between breakpoints if from intrachromosomal breakpoints, or intervals within a 100 kbp window on each side of interchromosomal breakpoints. The junction copy number (JCN) of the cluster is defined by the maximum junction copy number of any junction within the cluster. Low copy clusters thresholded at JCN < 4, and high copy junctions at JCN > 7. A junction-class dominant cluster is defined as a cluster containing at least 90% of all junctions of one of the 4 junction-classes. P-value obtained by Wilcoxon rank sum test. (B) Top, significance of enrichment within superenhancer peaks after correcting for covariates of replication timing. Q-Q plot of observed p-values illustrating the probability that the observed densities of pyrgo within superenhancers come from the expected gamma-Poisson binomial distribution and expected p-values from a uniform distribution. Labeling indicates that *MYC* falls within 500 kbp of the significantly pyrgo-enriched superenhancer. Bottom, fraction of cases that harbor pyrgo overlapping genes commonly amplified within GISTIC peaks from (Zack et al., 2013). Dots colored red in Q-Q plot are significant at *FDR* ≤ 0.10 threshold. (C) Top, significance of enrichment within genes after correcting for covariates of replication timing, gene length, and fragile site occurrence. Q-Q plot of observed p-values illustrating the probability that the observed densities of rigma within genes come from the expected gamma-Poisson distribution and expected p-values from a uniform distribution. Labeling indicates the top genes that are significantly enriched with rigma. Bottom, fraction of cases that harbor rigma overlapping with genes commonly deleted within GISTIC peaks from (Zack et al., 2013). (D) Fraction of cases harboring rigma within *FHIT* in specific gastric tumor types vs. all other tumor types (E) Representative patterns of pyrgo occurring near *MYC* and its associated superenhancer. (F) Representative pattern of rigma occurring within *FHIT* locus across multiple samples in the esophageal adenocarcinoma cohort (left) and the Barrett’s esophagus cohort (right). Abbreviations: CN, copy number; RD, read depth.

**Fig. S3.**
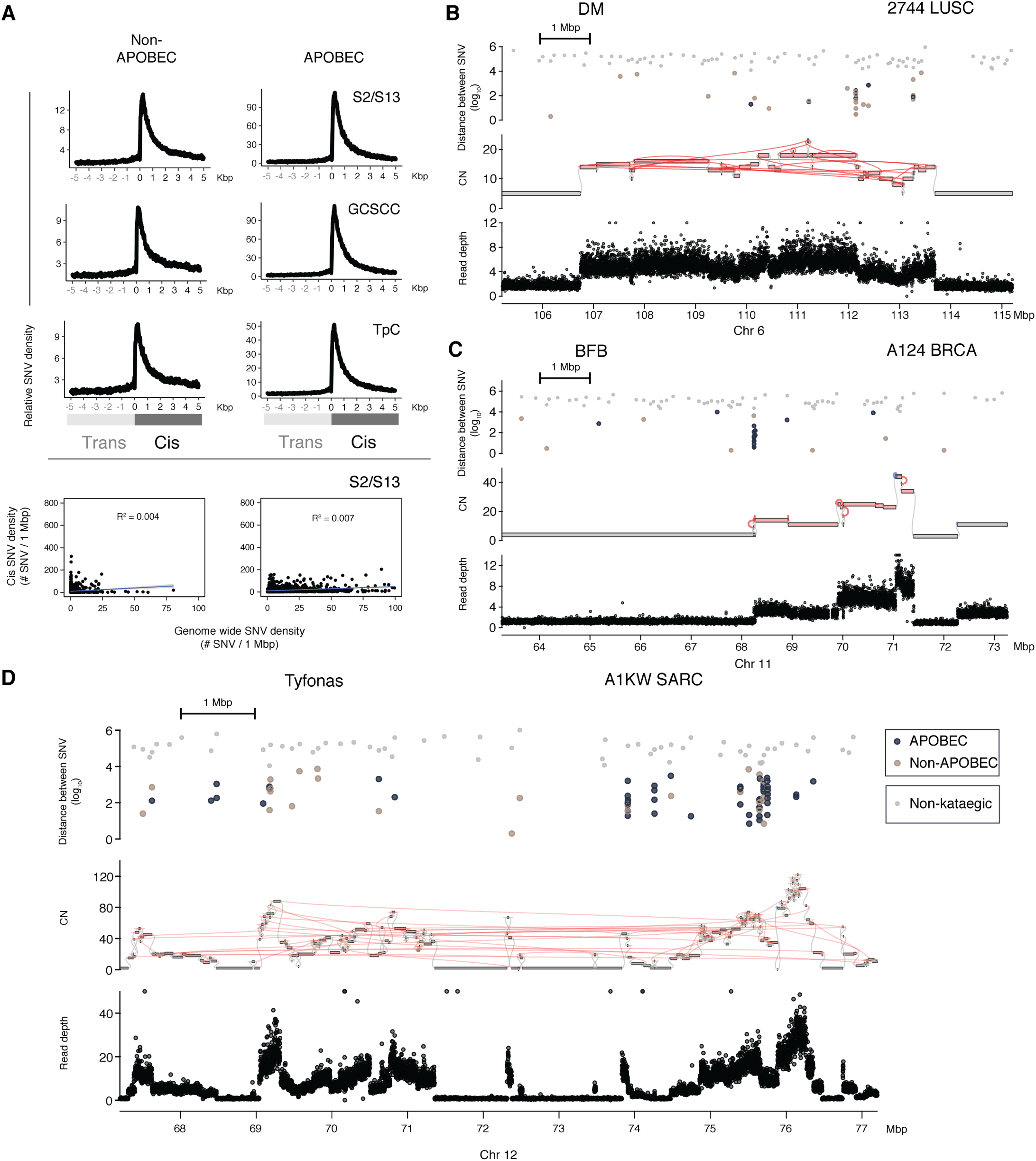
Kataegis occurs within and outside of APOBEC context defined by several methods. (A) Relative SNV densities on breakpoint centric coordinates shows an enrichment of SNV densities near junctions outside of APOBEC contexts (left column) as defined by COSMIC SNV signatures 2 and 13 (S2/S13), GC-strand coordinated clusters (described in ref. (Roberts et al., 2013)), or TpC mutational context (right column). Bottom panels, *cis* SNV density and genome wide SNV density are not correlated with one another, indicating the enrichment of SNV near junctions attributed to APOBEC- and non-APOBEC-related processes is independent of active processes genome-wide. Illustrations of kataegis by rainfall plots showing the inter SNV distance and among different event types: (B) tyfonas, (C) DM, and (D), BFBC. Dots are colored blue (APOBEC) and beige (non-APOBEC) for inter-SNV distances ≤ 1 × 10^4^ base pairs. Grey dots indicate SNV with inter-SNV of > 1 × 10^4^ base pairs. Inter-SNV distance is defined by distance of each SNV to the next on hg19 reference coordinates. Abbreviation: CN, copy number.

**Fig. S4.**
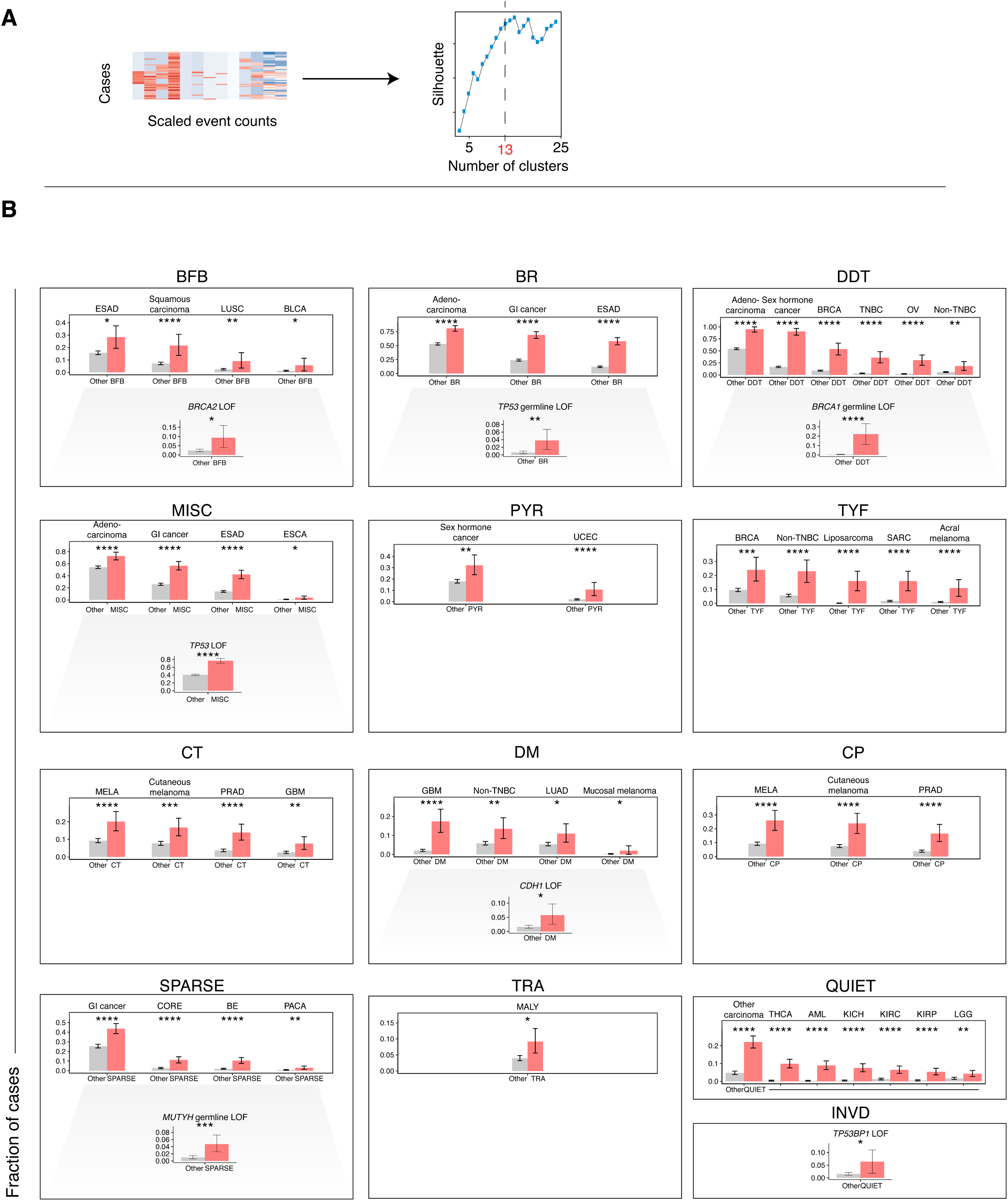
Clustering tuning and other tumor type and genotype associations in clusters. (A) Selecting number of clusters from event type features across 2,798 genome graphs. By elbow method, we visually identified 13 clusters as the most likely optimal number of clusters from our dataset using the Silhouette clustering metric. (B), each panel presents the significant tumor type enrichments for each of the 13 clusters identified from hierarchical clustering of the event type features among all cases in the cohort in **Fig. 5B-C**. Within each panel, the bottom row shows the significantly enriched genotypes within that cluster. Significance levels: **** (*FDR* < 1 × 10^−4^), *** (*FDR* < 1 × 10^−3^), ** (*FDR* < 0.01), * (*FDR* < 0.10).

**Fig. S5.**
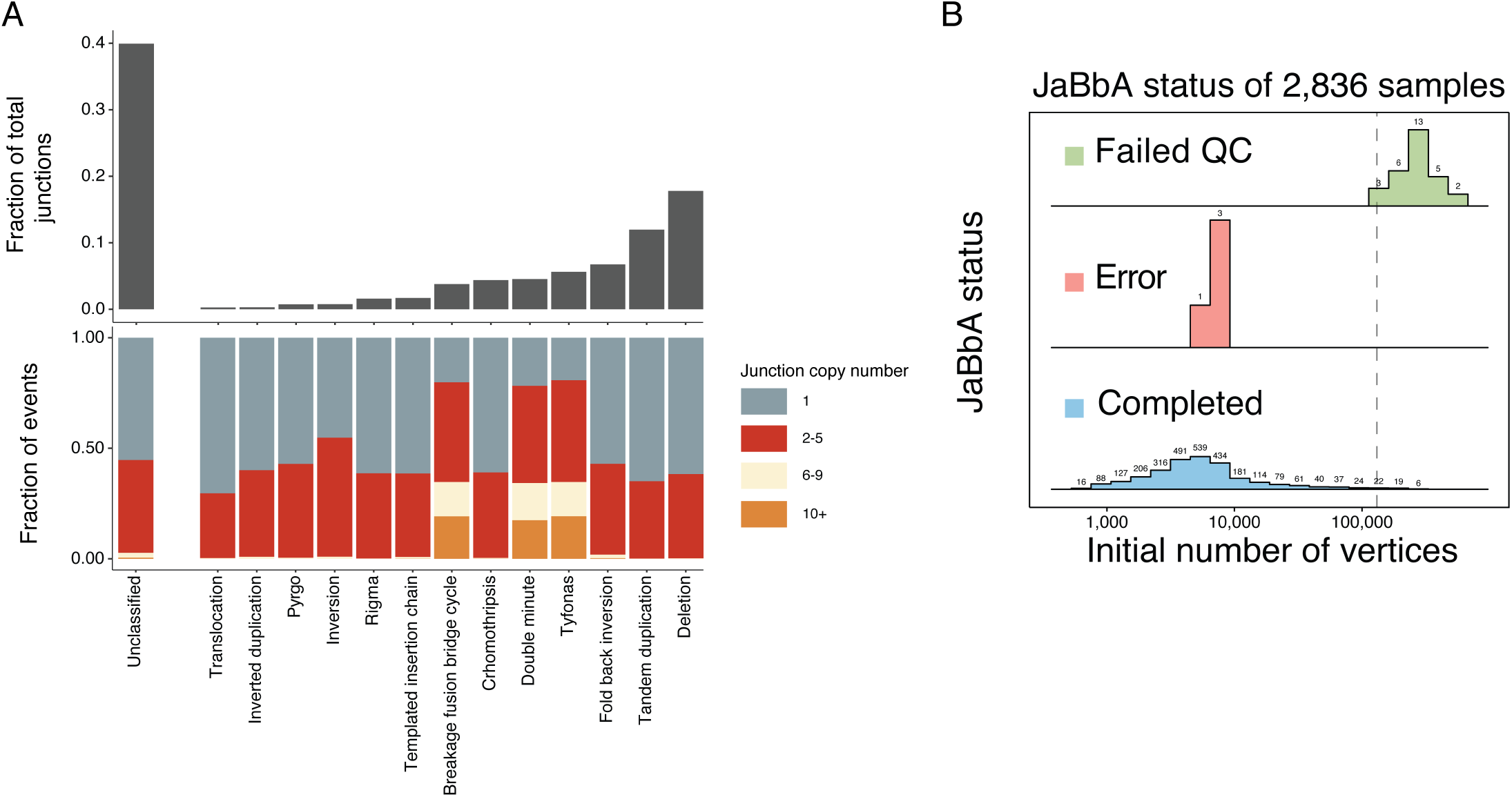
Completeness of event annotations and samples. (A) Unclassified rearrangements. Top, Fraction of the 308,137 junctions fitted in 2,798 genome graphs, that are classified within this article into event types. Bottom, fraction of each event type that falls within 4 different ranges of junction copy number as estimated by `JaBbA`. (B) 35 of the total 2,833 samples are not considered in the analysis due to various reasons. Bad quality samples are defined as larger than 130,000 initial total vertices to be optimized.

## Algorithms

### Genome graph construction

Algorithm S1 demonstrates how we build a genome graph from a set of adjacencies and genomic breakpoints (e.g. change-points inferred from analysis of binned read-depth data).

#### Algorithm S1 BuildGraph

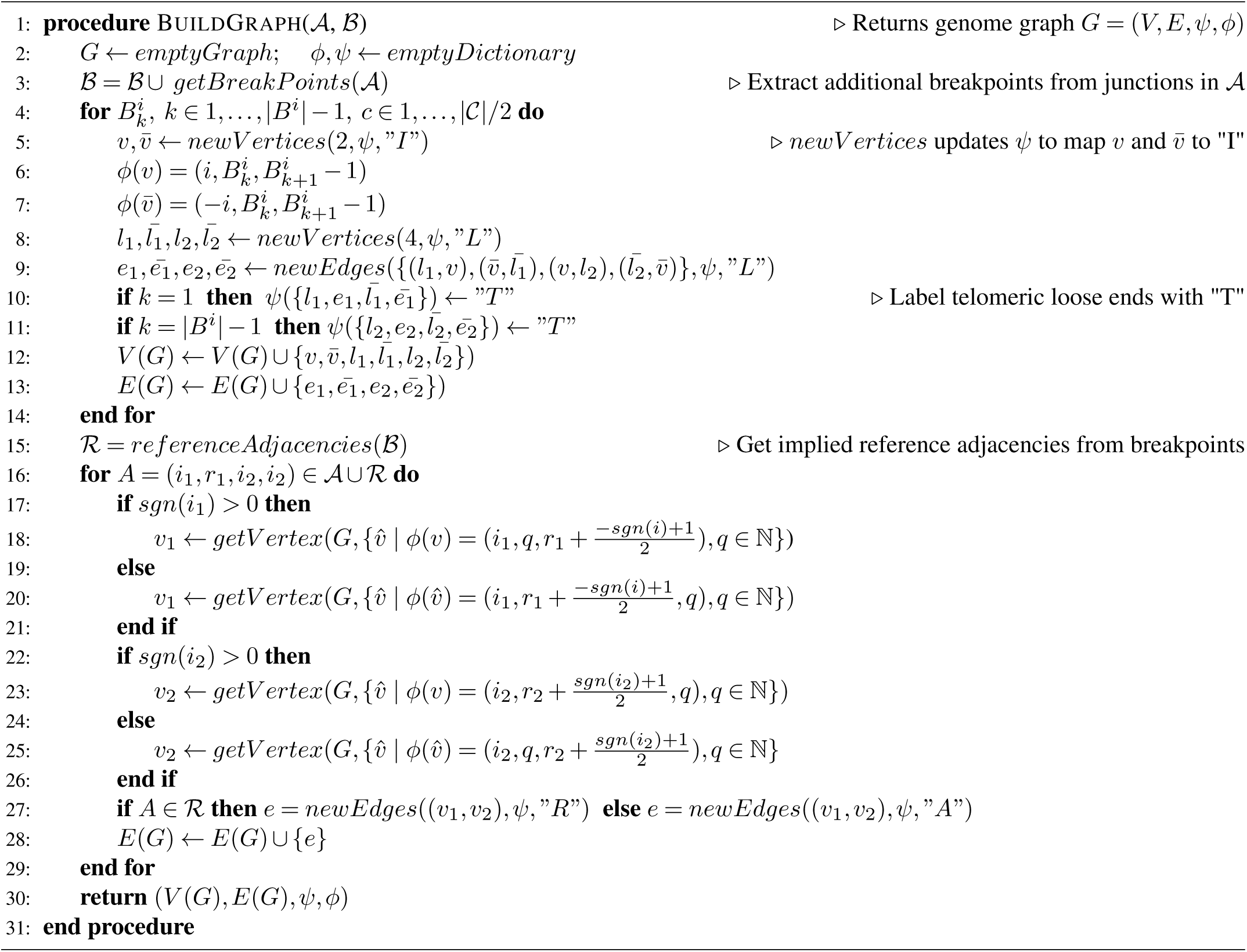

## References

Alexandrov, L. B., Nik-Zainal, S., Wedge, D. C., Aparicio, S. A. J. R., Behjati, S., Biankin, A. V., Bignell, G. R., Bolli, N., Borg, A., Børresen-Dale, A.-L., Boyault, S., Burkhardt, B., Butler, A. P., Caldas, C., Davies, H. R., Desmedt, C., Eils, R., Eyfjörd, J. E., Foekens, J. A., Greaves, M., Hosoda, F., Hutter, B., Ilicic, T., Imbeaud, S., Imielinski, M., Jäger, N., Jones, D. T. W., Jones, D., Knappskog, S., Kool, M., Lakhani, S. R., López-Otín, C., Martin, S., Munshi, N. C., Nakamura, H., Northcott, P. A., Pajic, M., Papaemmanuil, E., Paradiso, A., Pearson, J. V., Puente, X. S., Raine, K., Ramakrishna, M., Richardson, A. L., Richter, J., Rosenstiel, P., Schlesner, M., Schumacher, T. N., Span, P. N., Teague, J. W., Totoki, Y., Tutt, A. N. J., Valdés-Mas, R., Buuren, M. M. v., Veer, L. v., Vincent-Salomon, A., Waddell, N., Yates, L. R., Initiative, A. P. C. G., Consortium, I. B. C., Consortium, I. M.-S., PedBrain, I., Zucman-Rossi, J., Futreal, P. A., McDermott, U., Lichter, P., Meyerson, M., Grimmond, S. M., Siebert, R., Campo, E., Shibata, T., Pfister, S. M., Campbell, P. J. and Stratton, M. R. (2013). Signatures of mutational processes in human cancer. Nature.

Baca, S., Prandi, D., Lawrence, M., Mosquera, J., Romanel, A., Drier, Y., Park, K., Kitabayashi, N., MacDonald, T., Ghandi, M., Van Allen, E., Kryukov, G., Sboner, A., Theurillat, J.-P., Soong, T., Nickerson, E., Auclair, D., Tewari, A., Beltran, H., Onofrio, R., Boysen, G., Guiducci, C., Barbieri, C., Cibulskis, K., Sivachenko, A., Carter, S., Saksena, G., Voet, D., Ramos, A., Winckler, W., Cipicchio, M., Ardlie, K., Kantoff, P., Berger, M., Gabriel, S., Golub, T., Meyerson, M., Lander, E., Elemento, O., Getz, G., Demichelis, F., Rubin, M. and Garraway, L. (2013). Punctuated Evolution of Prostate Cancer Genomes. Cell.

Barretina, J., Caponigro, G., Stransky, N., Venkatesan, K., Margolin, A. A., Kim, S., Wilson, C. J., Lehár, J., Kryukov, G. V., Sonkin, D., Reddy, A., Liu, M., Murray, L., Berger, M. F., Monahan, J. E., Morais, P., Meltzer, J., Korejwa, A., Jané-Valbuena, J., Mapa, F. A., Thibault, J., Bric-Furlong, E., Raman, P., Shipway, A., Engels, I. H., Cheng, J., Yu, G. K., Yu, J., Aspesi, Jr, P., de Silva, M., Jagtap, K., Jones, M. D., Wang, L., Hatton, C., Palescandolo, E., Gupta, S., Mahan, S., Sougnez, C., Onofrio, R. C., Liefeld, T., MacConaill, L., Winckler, W., Reich, M., Li, N., Mesirov, J. P., Gabriel, S. B., Getz, G., Ardlie, K., Chan, V., Myer, V. E., Weber, B. L., Porter, J., Warmuth, M., Finan, P., Harris, J. L., Meyerson, M., Golub, T. R., Morrissey, M. P., Sellers, W. R., Schlegel, R. and Garraway, L. A. (2012). The Cancer Cell Line Encyclopedia enables predictive modelling of anticancer drug sensitivity. Nature.

Beroukhim, R., Mermel, C. H., Porter, D., Wei, G., Raychaudhuri, S., Donovan, J., Barretina, J., Boehm, J. S., Dobson, J., Urashima, M., Henry, K. T. M., Pinchback, R. M., Ligon, A. H., Cho, Y.-J., Haery, L., Greulich, H., Reich, M., Winckler, W., Lawrence, M. S., Weir, B. A., Tanaka, K. E., Chiang, D. Y., Bass, A. J., Loo, A., Hoffman, C., Prensner, J., Liefeld, T., Gao, Q., Yecies, D., Signoretti, S., Maher, E., Kaye, F. J., Sasaki, H., Tepper, J. E., Fletcher, J. A., Tabernero, J., Baselga, J., Tsao, M.-S., Demichelis, F., Rubin, M. A., Janne, P. A., Daly, M. J., Nucera, C., Levine, R. L., Ebert, B. L., Gabriel, S., Rustgi, A. K., Antonescu, C. R., Ladanyi, M., Letai, A., Garraway, L. A., Loda, M., Beer, D. G., True, L. D., Okamoto, A., Pomeroy, S. L., Singer, S., Golub, T. R., Lander, E. S., Getz, G., Sellers, W. R. and Meyerson, M. (2010). The landscape of somatic copy-number alteration across human cancers. Nature.

Bianchi, J. J., Murigneux, V., Bedora-Faure, M., Lescale, C. and Deriano, L. (2019). Breakage-Fusion-Bridge Events Trigger Complex Genome Rearrangements and Amplifications in Developmentally Arrested T Cell Lymphomas. Cell Reports.

Bignell, G. R., Greenman, C. D., Davies, H., Butler, A. P., Edkins, S., Andrews, J. M., Buck, G., Chen, L., Beare, D., Latimer, C., Widaa, S., Hinton, J., Fahey, C., Fu, B., Swamy, S., Dalgliesh, G. L., Teh, B. T., Deloukas, P., Yang, F., Campbell, P. J., Futreal, P. A. and Stratton, M. R. (2010). Signatures of mutation and selection in the cancer genome. Nature.

Boeva, V., Popova, T., Bleakley, K., Chiche, P., Cappo, J., Schleier-macher, G., Janoueix-Lerosey, I., Delattre, O. and Barillot, E. (2012). Control-FREEC: a tool for assessing copy number and allelic content using next-generation sequencing data. Bioinformatics.

Brunner, S. F., Roberts, N. D., Wylie, L. A., Moore, L., Aitken, S. J., Davies, S. E., Sanders, M. A., Ellis, P., Alder, C., Hooks, Y., Abascal, F., Stratton, M. R., Martincorena, I., Hoare, M. and Campbell, P. J. (2019). Somatic mutations and clonal dynamics in healthy and cirrhotic human liver. Nature.

Cameron, D. L., Schröder, J., Penington, J. S., Do, H., Molania, R., Dobrovic, A., Speed, T. P. and Papenfuss, A. T. (2017). GRIDSS: sensitive and specific genomic rearrangement detection using positional de Bruijn graph assembly. Genome Res.

Chen, X., Gupta, P., Wang, J., Nakitandwe, J., Roberts, K., Dalton, J. D., Parker, M., Patel, S., Holmfeldt, L., Payne, D., Easton, J., Ma, J., Rusch, M., Wu, G., Patel, A., Baker, S. J., Dyer, M. A., Shurtleff, S., Espy, S., Pounds, S., Downing, J. R., Ellison, D. W., Mullighan, C. G. and Zhang, J. (2015). CONSERTING: integrating copy-number analysis with structural-variation detection. Nat. Methods.

Cheng, J., Demeulemeester, J., Wedge, D. C., Vollan, H. K. M., Pitt, J. J., Russnes, H. G., Pandey, B. P., Nilsen, G., Nord, S., Bignell, G. R., White, K. P., Børresen-Dale, A.-L., Campbell, P. J., Kristensen, V. N., Stratton, M. R., Lingjærde, O. C., Moreau, Y. and Van Loo, P. (2017). Pan-cancer analysis of homozygous deletions in primary tumours uncovers rare tumour suppressors. Nat. Commun.

Chiang, D. Y., Getz, G., Jaffe, D. B., O’Kelly, M. J. T., Zhao, X., Carter, S. L., Russ, C., Nusbaum, C., Meyerson, M. and Lander, E. S. (2009). High-resolution mapping of copy-number alterations with massively parallel sequencing. Nature Methods.

Favero, F., Joshi, T., Marquard, A. M., Birkbak, N. J., Krzystanek, M., Li, Q., Szallasi, Z. and Eklund, A. C. (2015). Sequenza: allele-specific copy number and mutation profiles from tumor sequencing data. Ann. Oncol.

Frankell, A. M., Jammula, S., Li, X., Contino, G., Killcoyne, S., Abbas, S., Perner, J., Bower, L., Devonshire, G., Ococks, E., Grehan, N., Mok, J., O’Donovan, M., MacRae, S., Eldridge, M. D., Tavaré, S. and Fitzgerald, R. C. (2019). The landscape of selection in 551 esophageal adenocarcinomas defines genomic biomarkers for the clinic. Nature Genetics.

Fungtammasan, A., Walsh, E., Chiaromonte, F., Eckert, K. A. and Makova, K. D. (2012). A genome-wide analysis of common fragile sites: What features determine chromosomal instability in the human genome? Genome Research.

Funnell, T., Zhang, A. W., Grewal, D., McKinney, S., Bashashati, A., Wang, Y. K. and Shah, S. P. (2019). Integrated structural variation and point mutation signatures in cancer genomes using correlated topic models. PLOS Computational Biology.

Garsed, D., Marshall, O., Corbin, V., Hsu, A., Di Stefano, L., Schröder, J., Li, J., Feng, Z.-P., Kim, B., Kowarsky, M., Lansdell, B., Brookwell, R., Myklebost, O., Meza-Zepeda, L., Holloway, A., Pedeutour, F., Choo, K., Damore, M., Deans, A., Papenfuss, A. and Thomas, D. (2014). The Architecture and Evolution of Cancer Neochromosomes. Cancer Cell.

Ghandi, M., Huang, F. W., Jané-Valbuena, J., Kryukov, G. V., Lo, C. C., McDonald, E. R., Barretina, J., Gelfand, E. T., Bielski, C. M., Li, H., Hu, K., Andreev-Drakhlin, A. Y., Kim, J., Hess, J. M., Haas, B. J., Aguet, F., Weir, B. A., Rothberg, M. V., Paolella, B. R., Lawrence, M. S., Akbani, R., Lu, Y., Tiv, H. L., Gokhale, P. C., Weck, A. d., Mansour, A. A., Oh, C., Shih, J., Hadi, K., Rosen, Y., Bistline, J., Venkatesan, K., Reddy, A., Sonkin, D., Liu, M., Lehar, J., Korn, J. M., Porter, D. A., Jones, M. D., Golji, J., Caponigro, G., Taylor, J. E., Dunning, C. M., Creech, A. L., Warren, A. C., McFarland, J. M., Zamanighomi, M., Kauffmann, A., Stransky, N., Imielinski, M., Maruvka, Y. E., Cherniack, A. D., Tsherniak, A., Vazquez, F., Jaffe, J. D., Lane, A. A., Weinstock, D. M., Johannessen, C. M., Morrissey, M. P., Stegmeier, F., Schlegel, R., Hahn, W. C., Getz, G., Mills, G. B., Boehm, J. S., Golub, T. R., Garraway, L. A. and Sellers, W. R. (2019). Next-generation characterization of the Cancer Cell Line Encyclopedia. Nature.

Gisselsson, D., Pettersson, L., Höglund, M., Heidenblad, M., Gorunova, L., Wiegant, J., Mertens, F., Cin, P. D., Mitelman, F. and Mandahl, N. (2000). Chromosomal breakage-fusion-bridge events cause genetic intratumor heterogeneity. Proceedings of the National Academy of Sciences.

Greenman, C. D., Pleasance, E. D., Newman, S., Yang, F., Fu, B., Nik-Zainal, S., Jones, D., Lau, K. W., Carter, N., Edwards, P. A. W., Futreal, P. A., Stratton, M. R. and Campbell, P. J. (2012). Estimation of rearrangement phylogeny for cancer genomes. Genome Research.

Ha, G., Roth, A., Khattra, J., Ho, J., Yap, D., Prentice, L. M., Melnyk, N., McPherson, A., Bashashati, A., Laks, E., Biele, J., Ding, J., Le, A., Rosner, J., Shumansky, K., Marra, M. A., Gilks, C. B., Huntsman, D. G., McAlpine, J. N., Aparicio, S. and Shah, S. P. (2014). TITAN: inference of copy number architectures in clonal cell populations from tumor whole-genome sequence data. Genome Research.

Hayward, N. K., Wilmott, J. S., Waddell, N., Johansson, P. A., Field, M. A., Nones, K., Patch, A.-M., Kakavand, H., Alexandrov, L. B., Burke, H., Jakrot, V., Kazakoff, S., Holmes, O., Leonard, C., Sabarinathan, R., Mularoni, L., Wood, S., Xu, Q., Waddell, N., Tembe, V., Pupo, G. M., Paoli-Iseppi, R. D., Vilain, R. E., Shang, P., Lau, L. M. S., Dagg, R. A., Schramm, S.-J., Pritchard, A., Dutton-Regester, K., Newell, F., Fitzgerald, A., Shang, C. A., Grimmond, S. M., Pickett, H. A., Yang, J. Y., Stretch, J. R., Behren, A., Kefford, R. F., Hersey, P., Long, G. V., Cebon, J., Shackleton, M., Spillane, A. J., Saw, R. P. M., López-Bigas, N., Pearson, J. V., Thompson, J. F., Scolyer, R. A. and Mann, G. J. (2017). Whole-genome landscapes of major melanoma subtypes. Nature.

Hnisz, D., Abraham, B. J., Lee, T. I., Lau, A., Saint-André, V., Sigova, A. A., Hoke, H. A. and Young, R. A. (2013). Super-enhancers in the control of cell identity and disease. Cell.

Huang, W., Li, L., Myers, J. R. and Marth, G. T. (2012). ART: a next-generation sequencing read simulator. Bioinformatics.

Iliopoulos, D., Guler, G., Han, S.-Y., Druck, T., Ottey, M., Mc-Corkell, K. A. and Huebner, K. (2006). Roles of FHIT and WWOX fragile genes in cancer. Cancer Letters.

Imielinski, M., Guo, G. and Meyerson, M. (2017). Insertions and Deletions Target Lineage-Defining Genes in Human Cancers. Cell.

Kim, S., Scheffler, K., Halpern, A. L., Bekritsky, M. A., Noh, E., Källberg, M., Chen, X., Kim, Y., Beyter, D., Krusche, P. and Saunders, C. T. (2018). Strelka2: fast and accurate calling of germline and somatic variants. Nature Methods.

Korbel, J. and Campbell, P. (2013). Criteria for Inference of Chromothripsis in Cancer Genomes. Cell.

Lee, J. J.-K., Park, S., Park, H., Kim, S., Lee, J., Lee, J., Youk, J., Yi, K., An, Y., Park, I. K., Kang, C. H., Chung, D. H., Kim, T. M., Jeon, Y. K., Hong, D., Park, P. J., Ju, Y. S. and Kim, Y. T. (2019). Tracing Oncogene Rearrangements in the Mutational History of Lung Adenocarcinoma. Cell.

Lee-Six, H., Olafsson, S., Ellis, P., Osborne, R. J., Sanders, M. A., Moore, L., Georgakopoulos, N., Torrente, F., Noorani, A., Goddard, M., Robinson, P., Coorens, T. H. H., O’Neill, L., Alder, C., Wang, J., Fitzgerald, R. C., Zilbauer, M., Coleman, N., Saeb-Parsy, K., Martincorena, I., Campbell, P. J. and Stratton, M. R. (2019). The landscape of somatic mutation in normal colorectal epithelial cells. Nature.

Li, H. and Durbin, R. (2009). Fast and accurate short read alignment with Burrows-Wheeler transform. Bioinformatics.

Li, Y., Roberts, N., Weischenfeldt, J., Wala, J. A., Shapira, O., Schumacher, S., Khurana, E., Korbel, J. O., Imielinski, M., Beroukhim, R. and Campbell, P. (2017). Patterns of structural variation in human cancer. bioRxiv.

Li, Y., Zhou, S., Schwartz, D. C. and Ma, J. (2016). Allele-Specific Quantification of Structural Variations in Cancer Genomes. Cell Syst.

Maciejowski, J. and Imielinski, M. (2017). Modeling cancer rearrangement landscapes. Current Opinion in Systems Biology.

Macintyre, G., Goranova, T. E., Silva, D. D., Ennis, D., Piskorz, A. M., Eldridge, M., Sie, D., Lewsley, L.-A., Hanif, A., Wilson, C., Dowson, S., Glasspool, R. M., Lockley, M., Brockbank, E., Montes, A., Walther, A., Sundar, S., Edmondson, R., Hall, G. D., Clamp, A., Gourley, C., Hall, M., Fotopoulou, C., Gabra, H., Paul, J., Supernat, A., Millan, D., Hoyle, A., Bryson, G., Nourse, C., Mincarelli, L., Sanchez, L. N., Ylstra, B., Jimenez-Linan, M., Moore, L., Hofmann, O., Markowetz, F., McNeish, I. A. and Brenton, J. D. (2018). Copy number signatures and mutational processes in ovarian carcinoma. Nature genetics.

Mallick, S., Li, H., Lipson, M., Mathieson, I., Gymrek, M., Racimo, F., Zhao, M., Chennagiri, N., Nordenfelt, S., Tandon, A., Skoglund, P., Lazaridis, I., Sankararaman, S., Fu, Q., Rohland, N., Renaud, G., Erlich, Y., Willems, T., Gallo, C., Spence, J. P., Song, Y. S., Poletti, G., Balloux, F., van Driem, G., de Knijff, P., Romero, I. G., Jha, A. R., Behar, D. M., Bravi, C. M., Capelli, C., Hervig, T., Moreno-Estrada, A., Posukh, O. L., Balanovska, E., Balanovsky, O., Karachanak-Yankova, S., Sahakyan, H., Toncheva, D., Yepisko-posyan, L., Tyler-Smith, C., Xue, Y., Abdullah, M. S., Ruiz-Linares, A., Beall, C. M., Rienzo, A. D., Jeong, C., Starikovskaya, E. B., Metspalu, E., Parik, J., Villems, R., Henn, B. M., Hodoglugil, U., Mahley, R., Sajantila, A., Stamatoyannopoulos, G., Wee, J. T. S., Khusainova, R., Khusnutdinova, E., Litvinov, S., Ayodo, G., Comas, D., Hammer, M. F., Kivisild, T., Klitz, W., Winkler, C. A., Labuda, D., Bamshad, M., Jorde, L. B., Tishkoff, S. A., Watkins, W. S., Metspalu, M., Dryomov, S., Sukernik, R., Singh, L., Thangaraj, K., Pääbo, S., Kelso, J., Patterson, N. and Reich, D. (2016). The Simons Genome Diversity Project: 300 genomes from 142 diverse populations. Nature.

Martincorena, I., Fowler, J. C., Wabik, A., Lawson, A. R. J., Abascal, F., Hall, M. W. J., Cagan, A., Murai, K., Mahbubani, K., Stratton, M. R., Fitzgerald, R. C., Handford, P. A., Campbell, P. J., Saeb-Parsy, K. and Jones, P. H. (2018). Somatic mutant clones colonize the human esophagus with age. Science.

McPherson, A. W., Roth, A., Ha, G., Chauve, C., Steif, A., de Souza, C. P. E., Eirew, P., Bouchard-Côté, A., Aparicio, S., Sahinalp, S. C. and Shah, S. P. (2017). ReMixT: clone-specific genomic structure estimation in cancer. Genome Biol.

Medvedev, P., Fiume, M., Dzamba, M., Smith, T. and Brudno, M. (2010). Detecting copy number variation with mated short reads. Genome Research.

Menghi, F., Inaki, K., Woo, X., Kumar, P. A., Grzeda, K. R., Malhotra, A., Yadav, V., Kim, H., Marquez, E. J., Ucar, D., Shreckengast, P. T., Wagner, J. P., MacIntyre, G., Karuturi, K. R. M., Scully, R., Keck, J., Chuang, J. H. and Liu, E. T. (2016). The tandem duplicator phenotype as a distinct genomic configuration in cancer. Proceedings of the National Academy of Sciences.

Mrasek, K., Schoder, C., Teichmann, A.-C., Behr, K., Franze, B., Wilhelm, K., Blaurock, N., Claussen, U., Liehr, T. and Weise, A. (2010). Global screening and extended nomenclature for 230 aphidicolin-inducible fragile sites, including 61 yet unreported ones. International Journal of Oncology.

Nik-Zainal, S., Alexandrov, L. B., Wedge, D. C., Loo, P. V., Greenman, C. D., Raine, K., Jones, D., Hinton, J., Marshall, J., Stebbings, L. A., Menzies, A., Martin, S., Leung, K., Chen, L., Leroy, C., Ramakrishna, M., Rance, R., Lau, K. W., Mudie, L. J., Varela, I., McBride, D. J., Bignell, G. R., Cooke, S. L., Shlien, A., Gamble, J., Whitmore, I., Maddison, M., Tarpey, P. S., Davies, H. R., Papaemmanuil, E., Stephens, P. J., McLaren, S., Butler, A. P., Teague, J. W., Jönsson, G., Garber, J. E., Silver, D., Miron, P., Fatima, A., Boyault, S., Langerød, A., Tutt, A., Martens, J. W., Aparicio, S. A., Borg, A., Salomon, A. V., Thomas, G., Børresen-Dale, A.-L., Richardson, A. L., Neuberger, M. S., Futreal, P. A., Campbell, P. J., Stratton, M. R. and Consortium, t. B. C. W. G. o. t. I. C. G. (2012). Mutational Processes Molding the Genomes of 21 Breast Cancers. Cell.

Nik-Zainal, S., Davies, H., Staaf, J., Ramakrishna, M., Glodzik, D., Zou, X., Martincorena, I., Alexandrov, L. B., Martin, S., Wedge, D. C., Loo, P. V., Ju, Y. S., Smid, M., Brinkman, A. B., Morganella, S., Aure, M. R., Lingjærde, O. C., Langerød, A., Ringnér, M., Ahn, S.-M., Boyault, S., Brock, J. E., Broeks, A., Butler, A., Desmedt, C., Dirix, L., Dronov, S., Fatima, A., Foekens, J. A., Gerstung, M., Hooijer, G. K. J., Jang, S. J., Jones, D. R., Kim, H.-Y., King, T. A., Krishnamurthy, S., Lee, H. J., Lee, J.-Y., Li, Y., McLaren, S., Menzies, A., Mustonen, V., O’Meara, S., Pauporté, I., Pivot, X., Purdie, C. A., Raine, K., Ramakrishnan, K., Rodríguez-González, F. G., Romieu, G., Sieuwerts, A. M., Simpson, P. T., Shepherd, R., Stebbings, L., Stefansson, O. A., Teague, J., Tommasi, S., Treilleux, I., Eynden, G. G. V. d., Vermeulen, P., Vincent-Salomon, A., Yates, L., Caldas, C., Veer, L. v., Tutt, A., Knappskog, S., Tan, B. K. T., Jonkers, J., Borg, A., Ueno, N. T., Sotiriou, C., Viari, A., Futreal, P. A., Campbell, P. J., Span, P. N., Laere, S. V., Lakhani, S. R., Eyfjord, J. E., Thompson, A. M., Birney, E., Stunnenberg, H. G., Vijver, M. J. v. d., Martens, J. W. M., Børresen-Dale, A.-L., Richardson, A. L., Kong, G., Thomas, G. and Stratton, M. R. (2016). Landscape of somatic mutations in 560 breast cancer whole-genome sequences. Nature.

Oesper, L., Ritz, A., Aerni, S. J., Drebin, R. and Raphael, B. J. (2012). Reconstructing cancer genomes from paired-end sequencing data. BMC Bioinformatics.

Ohta, M., Inoue, H., Cotticelli, M. G., Kastury, K., Baffa, R., Palazzo, J., Siprashvili, Z., Mori, M., McCue, P., Druck, T., Croce, C. M. and Huebner, K. (1996). The FHIT Gene, Spanning the Chromosome 3p14.2 Fragile Site and Renal Carcinoma–Associated t(3;8) Breakpoint, Is Abnormal in Digestive Tract Cancers. Cell.

Olshen, A. B., Venkatraman, E. S., Lucito, R. and Wigler, M. (2004). Circular binary segmentation for the analysis of array-based DNA copy number data. Biostatistics.

Paulson, T. G., Galipeau, P. C., Oman, K. M., Sanchez, C. A., Kuhner, M. K., Smith, L. P., Shah, M., Arora, K., Shelton, J., Johnson, M., Corvelo, A., Maley, C. C., Yao, X., Hadi, K., Sanghvi, R., Venturini, E., Emde, A.-K., Hubert, B., Imielinski, M., Robine, N., Reid, B. J. and Li, X. (2019). Somatic genome dynamics of Barrett’s esophagus patients with non-cancer and cancer outcomes. In preparation.

Pearl, L. H., Schierz, A. C., Ward, S. E., Al-Lazikani, B. and Pearl, F. M. G. (2015). Therapeutic opportunities within the DNA damage response. Nature Reviews Cancer.

Reimann, J. D. R. and Fletcher, C. D. M. (2008). Chapter 37 - Soft- Tissue Sarcomas. In The Molecular Basis of Cancer (Third Edition), (Mendelsohn, J., Howley, P. M., Israel, M. A., Gray, J. W.and Thompson, C. B., eds), pp. 471–477. W.B. Saunders Philadelphia.

Roberts, S. A., Lawrence, M. S., Klimczak, L. J., Grimm, S. A., Fargo, D., Stojanov, P., Kiezun, A., Kryukov, G. V., Carter, S. L., Saksena, G., Harris, S., Shah, R. R., Resnick, M. A., Getz, G. and Gordenin, D. A. (2013). An APOBEC cytidine deaminase mutagenesis pattern is widespread in human cancers. Nature Genetics.

Rosenthal, R., McGranahan, N., Herrero, J., Taylor, B. S. and Swanton, C. (2016). deconstructSigs: delineating mutational processes in single tumors distinguishes DNA repair deficiencies and patterns of carcinoma evolution. Genome Biology.

Schwartz, M., Zlotorynski, E. and Kerem, B. (2006). The molecular basis of common and rare fragile sites. Cancer Letters.

Sedlazeck, F. J., Lee, H., Darby, C. A. and Schatz, M. C. (2018). Piercing the dark matter: bioinformatics of long-range sequencing and mapping. Nat. Rev. Genet.

Shen, R. and Seshan, V. E. (2016). FACETS: allele-specific copy number and clonal heterogeneity analysis tool for high-throughput DNA sequencing. Nucleic Acids Res.

Shoushtari, A. N., Munhoz, R. R., Kuk, D., Ott, P. A., Johnson, D. B., Tsai, K. K., Rapisuwon, S., Eroglu, Z., Sullivan, R. J., Luke, J. J., Gangadhar, T. C., Salama, A. K. S., Clark, V., Burias, C., Puzanov, I., Atkins, M. B., Algazi, A. P., Ribas, A., Wolchok, J. D. and Postow, M. A. (2016). The efficacy of anti-PD-1 agents in acral and mucosal melanoma. Cancer.

Sondka, Z., Bamford, S., Cole, C. G., Ward, S. A., Dunham, I. and Forbes, S. A. (2018). The COSMIC Cancer Gene Census: describing genetic dysfunction across all human cancers. Nat. Rev. Cancer.

Spies, N., Weng, Z., Bishara, A., McDaniel, J., Catoe, D., Zook, J. M., Salit, M., West, R. B., Batzoglou, S. and Sidow, A. (2017). Genome-wide reconstruction of complex structural variants using read clouds. Nature Methods.

Stephens, P. J., Greenman, C. D., Fu, B., Yang, F., Bignell, G. R., Mudie, L. J., Pleasance, E. D., Lau, K. W., Beare, D., Stebbings, L. A., McLaren, S., Lin, M.-L., McBride, D. J., Varela, I., NikZainal, S., Leroy, C., Jia, M., Menzies, A., Butler, A. P., Teague, J. W., Quail, M. A., Burton, J., Swerdlow, H., Carter, N. P., Mors-berger, L. A., Iacobuzio-Donahue, C., Follows, G. A., Green, A. R., Flanagan, A. M., Stratton, M. R., Futreal, P. A. and Campbell, P. J. (2011). Massive Genomic Rearrangement Acquired in a Single Catastrophic Event during Cancer Development. Cell.

Tate, J. G., Bamford, S., Jubb, H. C., Sondka, Z., Beare, D. M., Bindal, N., Boutselakis, H., Cole, C. G., Creatore, C., Dawson, E., Fish, P., Harsha, B., Hathaway, C., Jupe, S. C., Kok, C. Y., Noble, K., Ponting, L., Ramshaw, C. C., Rye, C. E., Speedy, H. E., Stefancsik, R., Thompson, S. L., Wang, S., Ward, S., Campbell, P. J. and Forbes, S. A. (2018). COSMIC: the Catalogue Of Somatic Mutations In Cancer. Nucleic Acids Research.

Verhaak, R. G. W., Bafna, V. and Mischel, P. S. (2019). Extra-chromosomal oncogene amplification in tumour pathogenesis and evolution. Nature Reviews Cancer.

Viswanathan, S. R., Ha, G., Hoff, A. M., Wala, J. A., Carrot-Zhang, J., Whelan, C. W., Haradhvala, N. J., Freeman, S. S., Reed, S. C., Rhoades, J., Polak, P., Cipicchio, M., Wankowicz, S. A., Wong, A., Kamath, T., Zhang, Z., Gydush, G. J., Rotem, D., Love, J. C., Getz, G., Gabriel, S., Zhang, C.-Z., Dehm, S. M., Nelson, P. S., Allen, E. M. V., Choudhury, A. D., Adalsteinsson, V. A., Beroukhim, R., Taplin, M.-E. and Meyerson, M. (2018). Structural Alterations Driving Castration-Resistant Prostate Cancer Revealed by Linked-Read Genome Sequencing. Cell.

Wala, J. A., Bandopadhayay, P., Greenwald, N., O’Rourke, R., Sharpe, T., Stewart, C., Schumacher, S., Li, Y., Weischenfeldt, J., Yao, X., Nusbaum, C., Campbell, P., Getz, G., Meyerson, M., Zhang, C.-Z., Imielinski, M. and Beroukhim, R. (2018). SvABA: genome-wide detection of structural variants and indels by local assembly. Genome research.

Willis, N. A., Frock, R. L., Menghi, F., Duffey, E. E., Panday, A., Camacho, V., Hasty, E. P., Liu, E. T., Alt, F. W. and Scully, R. (2017). Mechanism of tandem duplication formation in BRCA1-mutant cells. Nature.

Xi, R., Hadjipanayis, A. G., Luquette, L. J., Kim, T.-M., Lee, E., Zhang, J., Johnson, M. D., Muzny, D. M., Wheeler, D. A., Gibbs, R. A., Kucherlapati, R. and Park, P. J. (2011). Copy number variation detection in whole-genome sequencing data using the Bayesian information criterion. Proc. Natl. Acad. Sci. U. S. A.

Zack, T. I., Schumacher, S. E., Carter, S. L., Cherniack, A. D., Saksena, G., Tabak, B., Lawrence, M. S., Zhang, C.-Z., Wala, J., Mermel, C. H., Sougnez, C., Gabriel, S. B., Hernandez, B., Shen, H., Laird, P. W., Getz, G., Meyerson, M. and Beroukhim, R. (2013). Pan-cancer patterns of somatic copy number alteration. Nature Genetics.

Zakov, S., Kinsella, M. and Bafna, V. (2013). An algorithmic approach for breakage-fusion-bridge detection in tumor genomes. Proc. Natl. Acad. Sci. U. S. A.

